# A real data-driven simulation strategy to select an imputation method for mixed-type trait data

**DOI:** 10.1101/2022.05.03.490388

**Authors:** Jacqueline A. May, Zeny Feng, Sarah J. Adamowicz

## Abstract

Missing observations in trait datasets pose an obstacle for analyses in myriad biological disciplines. Considering the mixed results of imputation, the wide variety of available methods, and the varied structure of real trait datasets, a framework for selecting a suitable imputation method is advantageous. We invoked a real data-driven simulation strategy to select an imputation method for a given mixed-type (categorical, count, continuous) target dataset. Candidate methods included mean/mode imputation, *k*-nearest neighbour, random forests, and multivariate imputation by chained equations (MICE). Using a trait dataset of squamates (lizards and amphisbaenians; order: Squamata) as a target dataset, a complete-case dataset consisting of species with nearly complete information was formed for the imputation method selection. Missing data were induced by removing values from this dataset under different missingness mechanisms: missing completely at random (MCAR), missing at random (MAR), and missing not at random (MNAR). For each method, combinations with and without phylogenetic information from single gene (nuclear and mitochondrial) or multigene trees were used to impute the missing values for five numerical and two categorical traits. The performances of the methods were evaluated under each missing mechanism by determining the mean squared error and proportion falsely classified rates for numerical and categorical traits, respectively. A random forest method supplemented with a nuclear-derived phylogeny resulted in the lowest error rates for the majority of traits, and this method was used to impute missing values in the original dataset. Data with imputed values better reflected the characteristics and distributions of the original data compared to complete-case data. However, caution should be taken when imputing trait data as phylogeny did not always improve performance for every trait and in every scenario. Ultimately, these results support the use of a real data-driven simulation strategy for selecting a suitable imputation method for a given mixed-type trait dataset.

**Author summary:** The issue of missing data is problematic in trait datasets as the missingness pattern may not be entirely random. Whether data are missing may depend on other known observations in the dataset, or on the value of the missing data points themselves. When only complete cases are used in an analysis, derived results may be biased. Imputation is an alternative to complete-case analysis and entails filling in the missing values using information provided by other trait values present in the dataset. Including phylogenetic information in the imputation process can improve the accuracy of imputed values, though results are dependent on the amount and pattern of missingness. Most previous evaluations of imputation methods for trait datasets are limited to numerical simulated data, with categorical traits not considered. Given a particular dataset, we propose the use of a real data-driven simulation strategy to select an imputation method. We evaluated the accuracies of four different imputation methods, with and without phylogeny information, and under different simulated missingness patterns using an example reptile trait dataset. Results indicated that data imputed using the best-performing method better reflected the original dataset characteristics compared to complete-case data. As imputation performance varies depending on the properties of a given dataset, a real data-driven simulation strategy can be used to provide guidance on best imputation practices.

## Introduction

Trait data are used in a wide variety of biological disciplines, including evolutionary biology, community ecology, and biodiversity conservation. For instance, trait data pertaining to the life history of a species, such as longevity, metabolic rate, and generation time, are integral in studies of biological aging [1,2]. Environmental trait data, such as latitude, temperature, and habitat type, may be used to identify those species most at risk of extinction [3,4]. However, an extensive proportion of these trait data are often missing. Missingness may stem from a taxonomic bias: data are available in copious amounts for well-researched or charismatic species and are lacking for endangered species or those that inhabit remote environments (e.g. deep sea) [5–7]. Mammal and bird taxa tend to be well sampled, and data for a large and diverse array of traits are available for many groups [8,9]. However, regional and phylogenetic biases are common in trait data for groups such as reptiles and amphibians, and observations are largely limited to body size and habitat traits [9]. Species traits are often tied to evolutionary history, a concept referred to as phylogenetic signal [10]. Closely related species can share the characteristics that render them elusive or difficult to study (e.g. small body size), resulting in patchy or unreliable data for entire taxonomic clades [5,6,8]. Certain types of trait data may also be easier to quantify (e.g. morphometric data) as opposed to traits that require arduous or invasive data collection techniques (e.g. age or reproductive data) [11] (see Fig 1 for a visualization of missingness in reptiles). When trait datasets are used in biological studies, these biases can lead researchers to make erroneous conclusions about the data. Consequently, the development of approaches and guidelines for handling missing data is an important area of research that spans across multiple biological disciplines.

**Fig 1.**
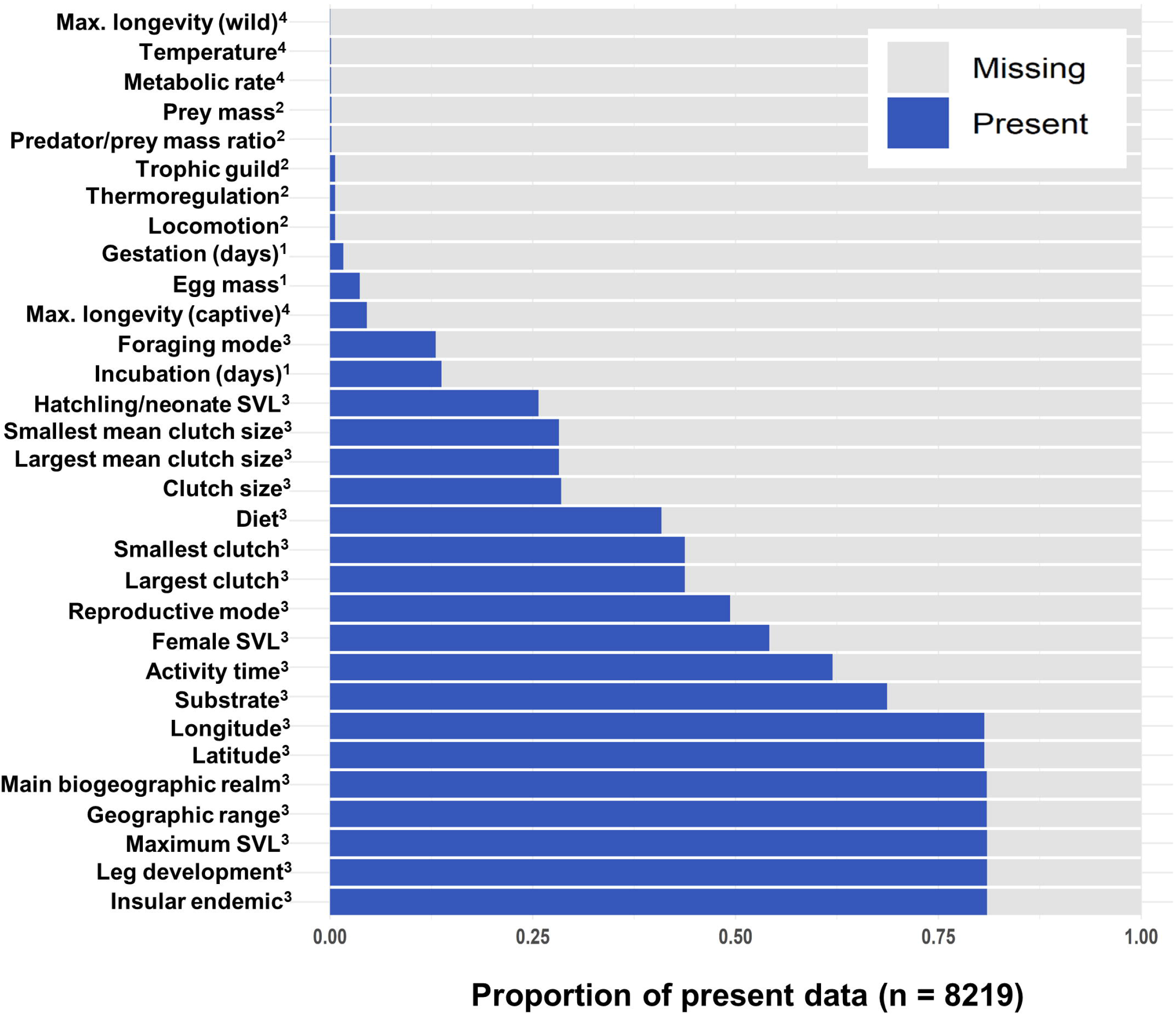
Visualization of missingness. Visualization of missingness (proportion of present vs. missing observations) in Squamata trait data obtained from the primary literature. Superscripts indicate the original sources of the trait data: 1) amniote life history database [12,13], 2) vertebrate home range sizes dataset [14,15], 3) traits of lizards of the world [16,17] and 4) AnAge [18,19]. See S1 File for further detail on trait sources.

The use of complete-case datasets can result in a large proportion of information being discarded [7,20]. More specifically, a loss in statistical power can result upon the removal of many incomplete cases if the data are “missing completely at random” (MCAR) [7,21]. Trait data, however, are often “missing at random” (MAR): observations that are missing for a particular trait are related to known values for other traits in the dataset. Simply removing incomplete cases when data are MAR can result in a reduction in statistical power and biased estimations of model parameters [7,11,22]. In more extreme cases, trait data may be “missing not at random” (MNAR): the reason data are missing is related to the unobserved data themselves. In such scenarios, the reason for missingness may be unclear to the researcher and thus difficult to verify empirically [23].

Imputing missing observations is a common alternative to the complete-case analysis. Imputation techniques use known observations to estimate the missing and unobserved values of a variable (or variables) of interest. Single imputation techniques such as hot deck imputation or *k*-nearest neighbour (KNN) [24] offer an efficient means for estimating missing values; however, these methods provide only a single estimate of the missing value. Random forest methods such as missForest [25] are also growing in popularity as they make no prior assumptions about the distributions of variables. Multiple imputation techniques have been developed that perform single imputation several times and are therefore capable of providing a measure of uncertainty of the imputed values [7,26]. An example of a multiple imputation method is multivariate imputation by chained equations (MICE; 27], which offers numerous models for imputing data of different types. Incorporating phylogenetic information into the imputation process has also been shown to increase the accuracy of imputed values [11,28]. This increase in accuracy is a result of the phylogenetic signal that is often inherent in trait data. A commonly used method for incorporating phylogenetic information into the imputation process is the use of phylogenetic eigenvectors. More specifically, methods such as phylogenetic eigenvector regression (PVR) [29] and phylogenetic eigenvector mapping (PEM) [30] employ a principal coordinates analysis (PCoA) to derive eigenvectors from a phylogenetic tree. PEM expands on the PVR method by applying an additional branch length transformation based on the Ornstein-Uhlenbeck evolutionary model [30,31]. Phylogenetic eigenvectors may then be used as additional predictor variables in the imputation process [11,32,33].

As missing data are a major concern in trait datasets, we are motivated to consider imputing these values. The correlative nature of trait data makes them suitable candidates for imputation, particularly when phylogenetic signal is also present [34]. When phylogenetic signal is strong for a trait, as may be the case for recently diverged groups, species may be more likely to exhibit similar values of the trait; these relationships can then be utilized by imputation methods to improve estimates. In an evaluation of imputation methods using mammalian trait data, Penone *et al*. [11] found that supplementing the imputation process with phylogenetic information improved the accuracy of KNN, missForest, and MICE for several life history traits. Kim *et al*. [32] similarly found that adding phylogenetic information to MICE improved accuracy rates of estimated functional diversity metrics. However, when imputing bird demographic traits with moderate phylogenetic signal (Pagel’s λ < 0.8), James *et al*. [35] found that use of phylogenetic information improved error rates by a margin of less than 1%. Moreover, they suggest that the use of auxiliary traits (traits that are present in the dataset but not the target of imputation) were often sufficient for accurate imputations. In sum, these findings indicate that improvements conferred by phylogenetic imputation methods are context-dependent, contingent upon the presence of phylogenetic signal and relationships among traits in the dataset.

Trait data exist in several forms, ranging from the discrete categories of foraging behaviour to the countable number of eggs in a nest. Many contemporary imputation methods are able to estimate mixed-type data, including both categorical and numerical (continuous and count) values. However, several previous studies have only evaluated their performances using simulated trait data [32,34], and the studies that have utilized real data are limited to numerical traits [11,36]. Given the diversity of available trait datasets and the varied success of imputation methods in comparative evaluations using trait data [11,34], additional guidelines are needed regarding imputation method selection. Identifying an imputation method that is most suitable for a given dataset, including its combinations of variables and biological properties, would be advantageous in reducing error rates when imputing missing trait values. Here, we propose a real data-driven strategy to select an imputation method for a given target trait dataset. The strategy entails four main steps: 1) missingness simulations under MCAR, MAR, and MNAR mechanisms, 2) imputation of simulated missing values using different candidate methods, 3) identification of the best-suited imputation method based on their performances, and 4) application of the best-suited imputation method to the target dataset and comparison between original, imputed, and complete-case data characteristics. The best-suited imputation method for the dataset is defined as the method, with or without phylogeny, that results in the lowest imputation error rates overall for both numerical and categorical traits. The real data simulation results can provide additional insights about the dataset in question including 1) whether there is an imputation method that performs best (i.e. results in the lowest imputation error rates) for a specific data type (continuous, count, categorical) and/or missingness mechanism (MCAR, MAR, and MNAR); 2) whether phylogenetic information improves the imputation performance; and 3) which type of phylogenetic information is influential (mitochondrial, nuclear). Phylogenetic resolution varies among gene trees [37,38], and certain genes may be more or less suited for imputation in a given taxon and taxonomic rank. Together, these insights can help determine a target dataset’s suitability for imputation including whether phylogenetic imputation is useful and if some traits are more amenable to imputation than others. The method-selection strategy proposed here may be considered for future trait-based analyses to avert biases associated with the use of complete case and to mitigate error rates by employing a well-suited imputation method. In turn, this can facilitate the pursuit of new research directions, particularly in those fields impeded by patchy and incomplete trait data.

## Results

### Performance comparison without phylogeny

Imputation performance was measured using mean squared error (MSE) and proportion falsely classified (PFC) for numerical and categorical traits, respectively. In both cases, lower error rates are indicative of better performance. In general, when missing data were simulated under MCAR, error rate increased with missingness proportion; this trend was observed for all trait and method combinations (Fig 2). Under the same simulation settings, *k*-nearest neighbour (*KNN)* [24,39], random forests (*RF*; “missForest” R package) [25,40] and multivariate imputation by chained equations (*MICE)* [27] outperformed mode and mean imputation for the majority of traits. However, for the multi-level categorical trait activity time, mode imputation resulted in a lower error rate than *RF* and *KNN* at 30-40% missingness (Fig 2a). Additionally, for smallest clutch, the mean imputation method outperformed *KNN* (10-40%) and *RF* (10%, 30-40%) (Fig 2d). *MICE* resulted in lower error rates than *KNN* and *RF* for most traits across all missingness proportions. However, *KNN* resulted in the lowest error rate for activity time at 10% missingness, and *RF* resulted in the lowest error rate across all missingness proportion settings for the insular endemic trait (Fig 2b). Under MAR without phylogeny, *MICE* generally outperformed both *RF* and *KNN* (see Figs 3 and 4); however, results were mixed for MNAR scenarios without phylogeny as the prevailing method depended on the trait in question. For example, *KNN* performed best for smallest clutch under MNAR without phylogeny (Fig 4b) whereas *RF* performed best for both categorical traits (Fig 3).

**Fig 2.**
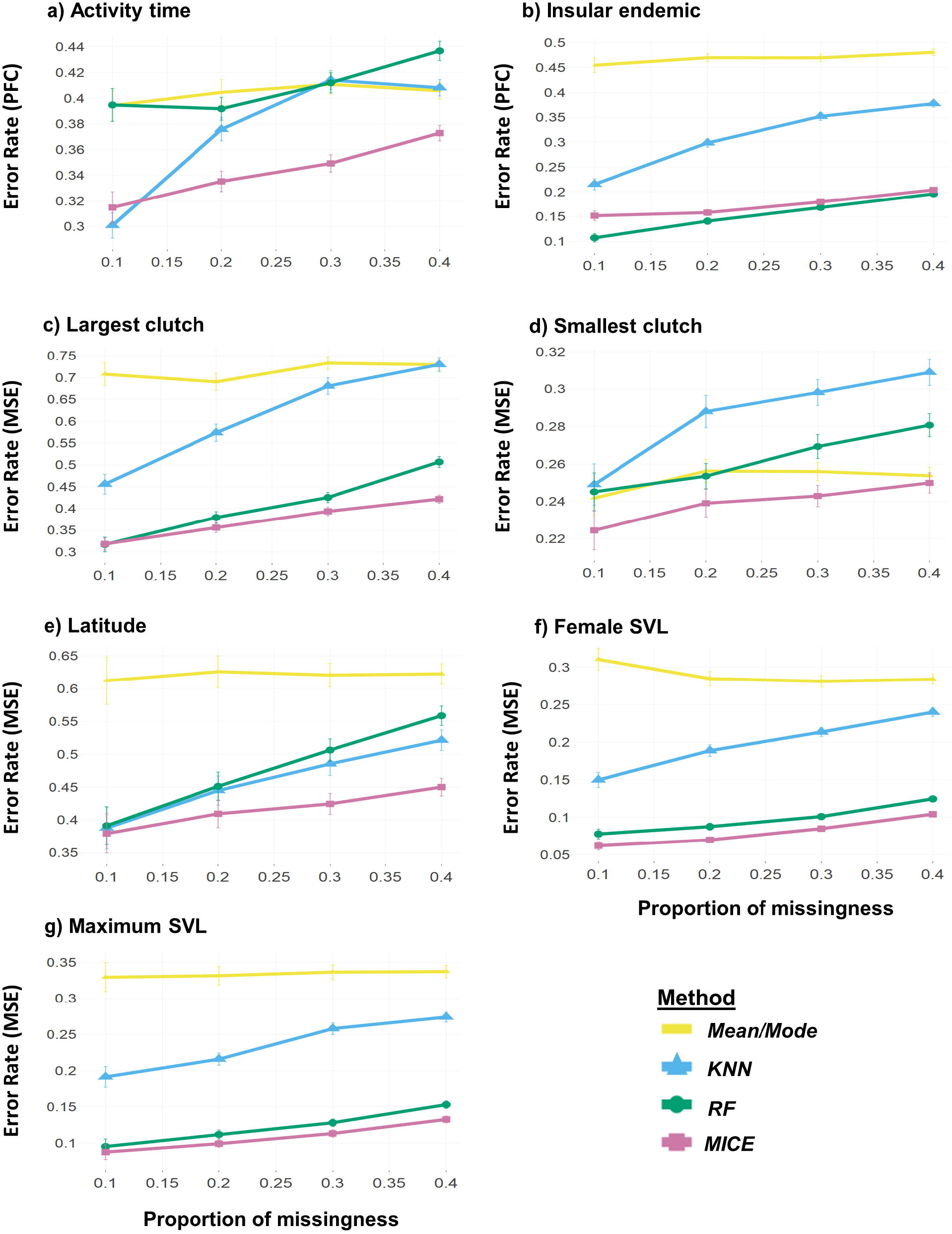
MCAR performance without phylogeny. Performances of the methods mean/mode imputation, *KNN*, missForest (*RF*), and multivariate imputation by chained equations (*MICE*) across different proportions of missingness when data were missing completely at random (MCAR). For MICE, logistic regression and predictive mean matching were used for imputing categorical and numerical traits, respectively. Error rate was measured as proportion falsely classified (PFC) for the categorical traits a) activity time and b) insular endemic and as mean squared error (MSE) for the numerical traits c) largest clutch, d) smallest clutch, e) latitude, f) female snout-vent length (SVL), and g) maximum SVL. PFC and MSE values for each trait were averaged across 100 dataset replicates. In both cases, error rates closer to 0 are indicative of better performance.

**Fig 3.**
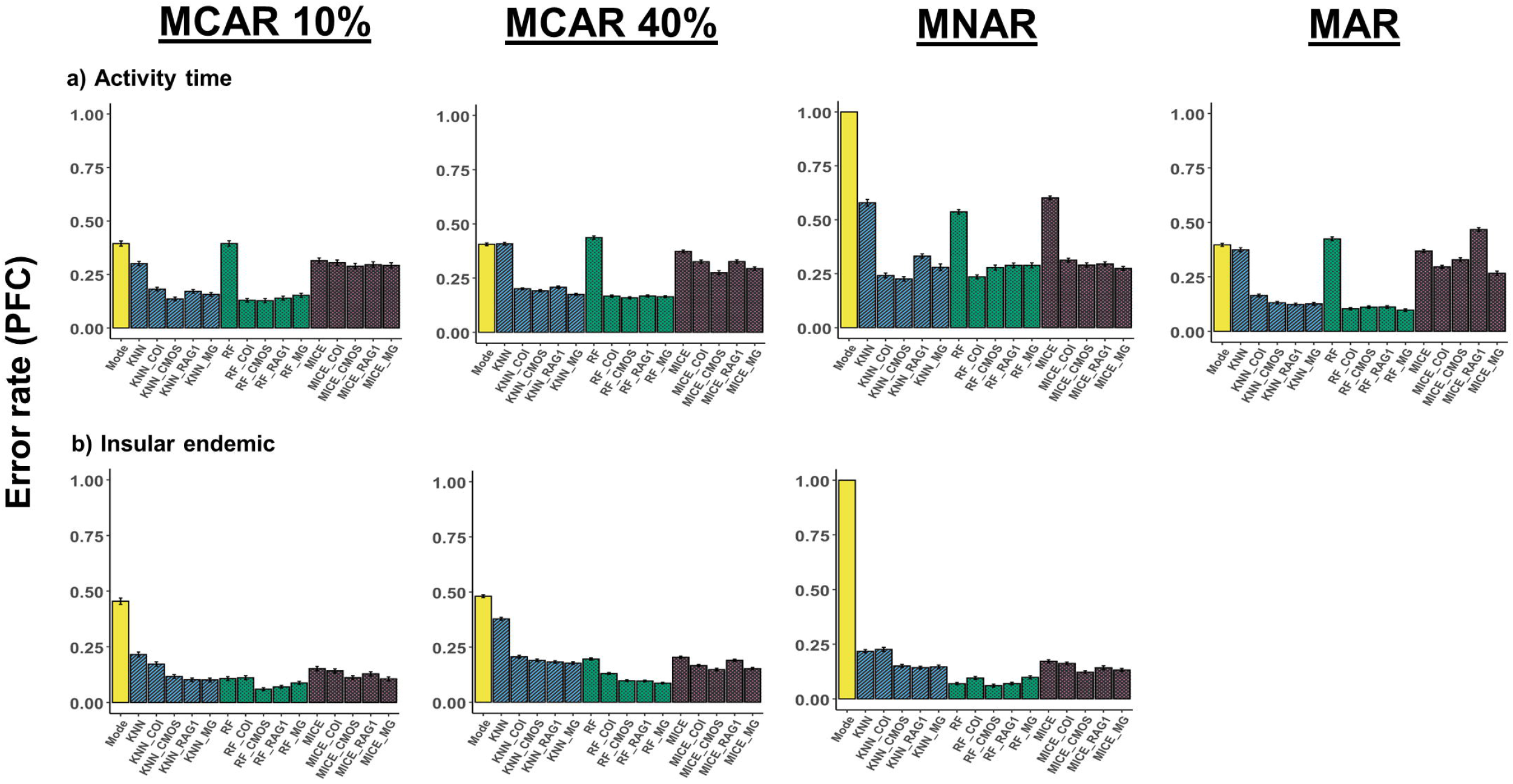
Imputation performance under different missingness mechanisms for categorical traits. Comparison of error rates (PFC) for the methods mean/mode imputation, *KNN, RF*, and *MICE* under different missingness mechanisms with and without the addition of phylogenetic information. Phylogenetic information was added in the form of trees built from sequence data of mitochondrial cytochrome *c* oxidase subunit I (COI), nuclear oocyte maturation factor (c-mos), recombination activating gene 1 (RAG1), and a composite multigene (MG) tree. Performances were quantified using PFC for the categorical traits a) activity time and b) insular endemic. PFC values for each trait were averaged across 100 dataset replicates. MAR = missing at random; MNAR = missing not at random.

**Fig 4.**
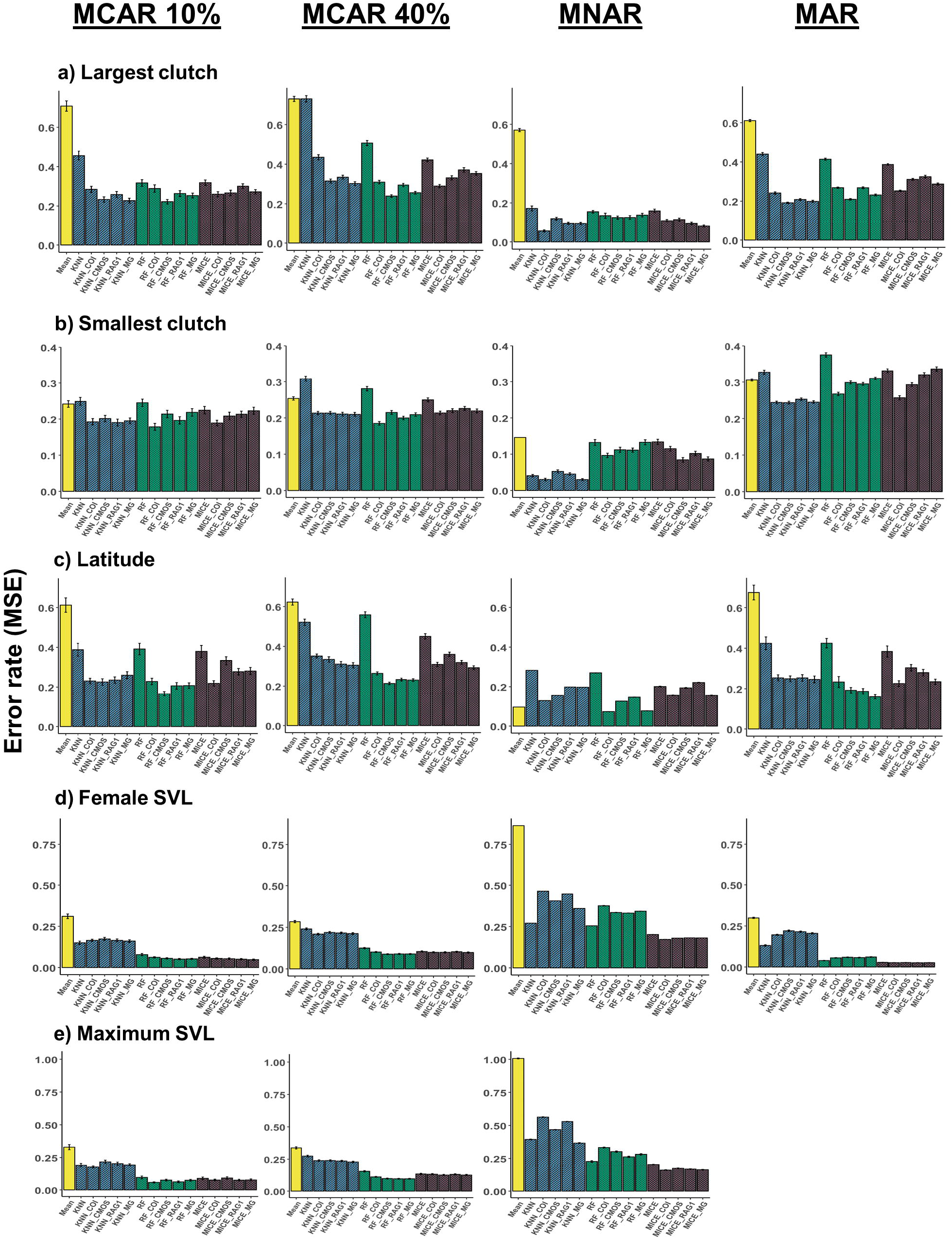
Imputation performance under different missingness mechanisms for numerical traits. Comparison of error rates for the methods mean/mode imputation, *KNN, RF*, and *MICE* under different missingness mechanisms with and without the addition of phylogenetic information. Phylogenetic information was added in the form of trees built from sequence data of COI, c-mos, RAG1, and a composite multigene (MG) tree. Performances were quantified using MSE for the numerical traits a) largest clutch, b) smallest clutch, c) latitude, f) female SVL, and g) maximum SVL. MSE values for each trait were averaged across 100 dataset replicates.

### Phylogenetic imputation performance

All traits exhibited significant phylogenetic signal in all gene trees (S1 Fig; see S1 File for more details on phylogenetic signal measures), and phylogenetic information was incorporated into the imputation process using phylogenetic eigenvectors (see SI). Although adding phylogeny was generally beneficial, improvements to imputation performance were contingent on method, data type, and missingness mechanism (Figs 3 and 4). For instance, when considering the categorical traits activity time and insular endemic, supplementing phylogenetic information from any of the trees generally improved performance for each method and under each missingness mechanism (Fig 3). Under the MAR scenario for activity time, however, adding RAG1 information to *MICE* resulted in an increased error rate relative to the scenario without phylogeny (Fig 3a). For most traits, MAR results reflected those under the MCAR mechanism; however, deviations from the general pattern occurred in some MNAR cases. For example, under MNAR for female SVL and maximum SVL, adding information from any tree increased error rates for both *KNN* and *RF* (Fig 4d,e).

For the traits largest clutch, smallest clutch, and latitude, *KNN, RF*, and *MICE* performances were improved (or remained neutral) after the addition of any type of phylogenetic information under MCAR, MAR, and MNAR (Fig 4a-c). However, which tree resulted in the best imputation performance varied depending on the trait, method, and missingness mechanism. For example, *RF* with c-mos performed best for largest clutch under MCAR 40% (Fig 4a), whereas *KNN* with COI or multigene (“MG”) performed best for smallest clutch under MNAR (Fig 4b). The traits female snout-vent length (SVL) and maximum SVL displayed somewhat dissimilar patterns from the other traits (Figs 4d,e) as phylogenetic information decreased imputation performance or had no substantial effect in most scenarios.

The relationship between phylogenetic signal and error ratio (error rate without phylogeny/error rate with phylogeny) varied depending on data type. For categorical traits, higher error ratio, indicative of better performance due to phylogeny, was generally associated with higher phylogenetic signal strength (Fig 5a). This same pattern was not as apparent for numerical traits (Fig 5b). Moreover, under the MNAR mechanism for numerical traits, several error ratio values fell below 1 with stronger phylogenetic signal, indicative of a reduction in performance due to phylogeny.

**Fig 5.**
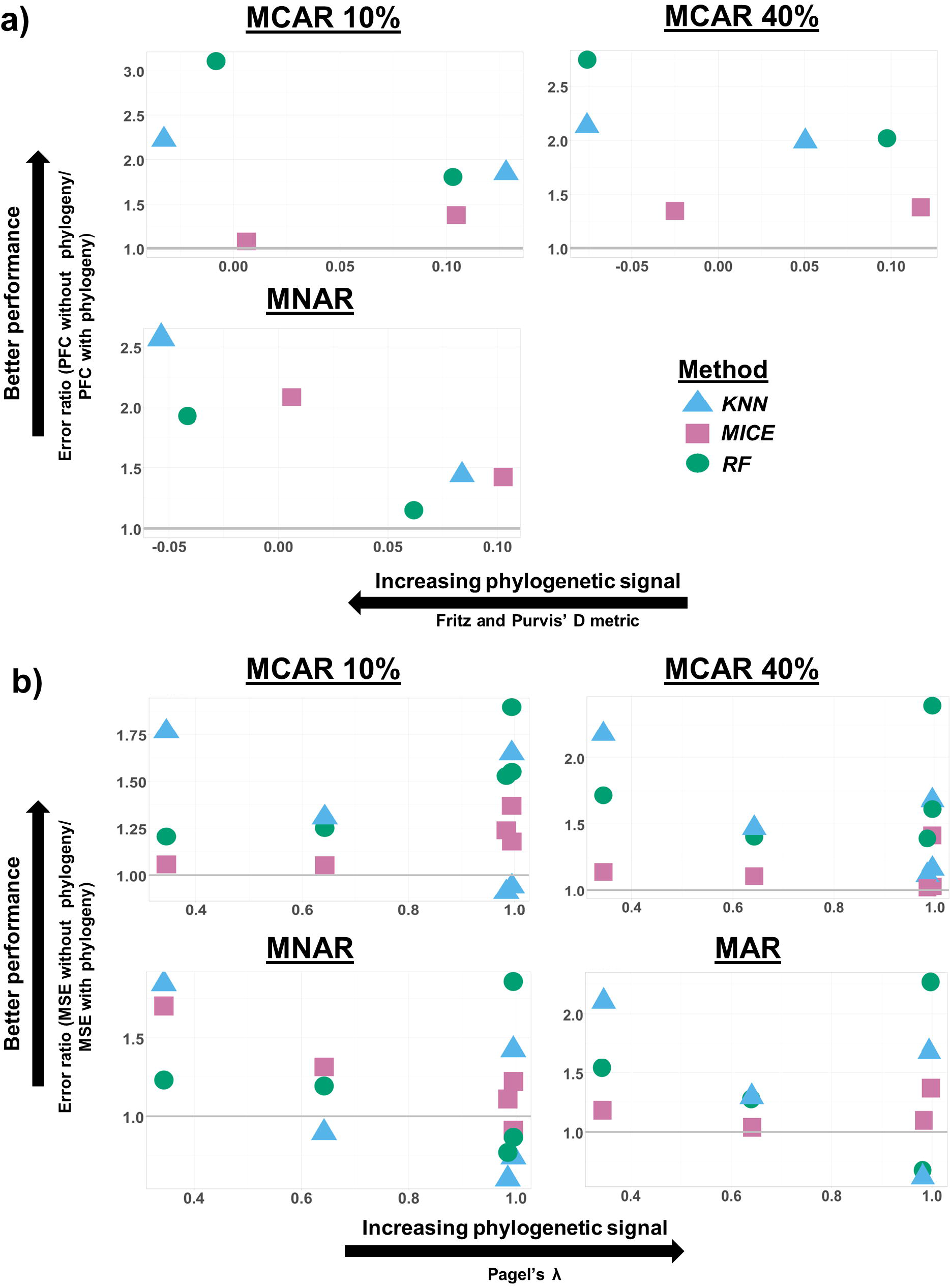
Association between error ratio and phylogenetic signal. Association between error ratio (error rate without phylogeny/error rate with phylogeny) and phylogenetic signal (Fritz and Purvis’ *D* [41] for categorical traits and Pagel’s λ [42] for numerical traits) at different proportions of missingness. Error ratio values above 1 (indicated by the gray line) signify an improvement in performance when phylogeny is added. In the case of *D*, lower values are indicative of higher levels of phylogenetic conservation for the trait; conversely, higher values of λ suggest stronger phylogenetic signal. Results are not shown for MAR in a) as only one trait (activity time) was simulated under this mechanism. To improve visualization, values were jittered (random noise introduced to data) using the package “ggplot2” [43]. Additionally, results are only shown for the c-mos gene (categorical traits) and RAG1 gene (numerical traits) as other genes followed similar patterns and these genes displayed the greatest range in phylogenetic signal.

### Imputation of original dataset using best-suited method

Using our majority voting system, the method that resulted in the lowest error rates for the majority of traits was *RF* with c-mos. Consequently, *RF* with c-mos was considered as the best-suited method for this dataset and used to impute the missing values in the target trait dataset (hereafter, “original dataset”). Out of the total species in the original dataset (*n* = 6657), those with available information for any of the seven traits as well as available c-mos sequence records were included in the subset of original data to be imputed (*n* = 921). The proportion of missingness varied for each trait in this original data subset as 0.16 for activity time, 0 for insular endemic, 0.21 for largest clutch, 0.21 for smallest clutch, 0.23 for female SVL, 0 for maximum SVL, and 0 for latitude. As insular endemic, maximum SVL, and latitude had complete observations in this subset, these traits did not require imputation.

Distributions and categorical frequencies of the complete-case, original, and original data with imputed values (hereafter, “imputed data”) can be observed in Fig 6. For the trait activity time, when compared to the original data, discrepancies in the categorical frequencies were more apparent in the complete-case data than in the imputed data (Fig 6a). The complete-case data displayed a greater overrepresentation of the rarest category (cathemeral: 11% vs. 8.9%) and underrepresentation of the most common category (diurnal: 57.9% vs. 64.4%). Conversely, the imputed data displayed a greater representation of observations in the most common category compared to the original data (diurnal: 67.5% vs. 64.4%). For all numerical traits, the imputed data distributions followed the distributions of the original more closely than did the complete-case distributions (Fig 6b-d; Table 1). Perhaps most apparent are the discrepancies in the maximum values in the complete-case data compared to those in the original and imputed data (e.g. for largest clutch, 68 vs. 88; for smallest clutch, 8 vs. 30). Although both complete-case and imputed data displayed reduced variance relative to the original data for the traits largest clutch, smallest clutch, and female SVL, the loss in variance for the complete-case data was considerably greater for all traits.

**Table 1.** Summary statistics for the complete-case data, original data, and original data with imputed values.

|  | Largest clutch<br>(# eggs/neonates) |  |  | Smallest clutch<br>(# eggs/neonates) |  |  | Female SVL<br>(mm) |  |  | Maximum SVL<br>(mm) |  | Latitude (°) |  |
| --- | --- | --- | --- | --- | --- | --- | --- | --- | --- | --- | --- | --- | --- |
|  | *CC* | *O* | *I* | *CC* | *O* | *I* | *CC* | *O* | *I* | *CC* | *O* | *CC* | *O* |
| **N** | 141 | 731 | 921 | 141 | 731 | 921 | 137 | 705 | 921 | 152 | 921 | 152 | 921 |
| **Min** | 1 | 1 | 1 | 1 | 1 | 1 | 18.7 | 18.7 | 18.7 | 21.7 | 21.7 | -40.36 | -47.89 |
| **Max** | 68 | 88 | 88 | 8 | 30 | 30 | 499.5 | 534.3 | 534.3 | 1170 | 1170 | 56.6 | 56.6 |
| **Range** | 67 | 87 | 87 | 7 | 29 | 29 | 480.8 | 515.6 | 515.6 | 1148.3 | 1148.3 | 96.96 | 104.49 |
| **Median** | 2 | 3 | 3 | 1 | 2 | 2 | 60.1 | 62.7 | 65.2 | 77 | 80 | -11.36 | -9.48 |
| **Mean** | 5.79 | 6.08 | 6.06 | 1.63 | 2.06 | 2.14 | 75.82 | 83.4 | 84.36 | 103.44 | 110.35 | 1.65 | -3.8 |
| **SE (mean)** | 0.67 | 0.33 | 0.27 | 0.09 | 0.07 | 0.06 | 4.76 | 2.43 | 2.07 | 8.58 | 3.30 | 2.03 | 0.75 |
| **0.95 CI (mean)** | 1.33 | 0.65 | 0.53 | 0.18 | 0.14 | 0.12 | 9.41 | 4.76 | 4.06 | 16.96 | 6.48 | 4.01 | 1.48 |
| **Variance** | 63.85 | 79.46 | 67.41 | 1.22 | 3.90 | 3.46 | 3103.49 | 4151.01 | 3950.16 | 11197.05 | 10048.39 | 627.38 | 521.53 |
| **Standard deviation** | 7.99 | 8.91 | 8.21 | 1.10 | 1.98 | 1.86 | 55.71 | 64.43 | 62.85 | 105.82 | 100.24 | 25.05 | 22.84 |

Summary statistics of the complete-case (CC), original (O), and original data with imputed values (I) for the numerical traits largest clutch, smallest clutch, female snout-vent length (SVL), and latitude. As the proportion of missingness was 0 for the traits maximum SVL and latitude in the original data subset, these traits were not imputed. Original trait data obtained from Meiri [16,17].

**Fig 6.**
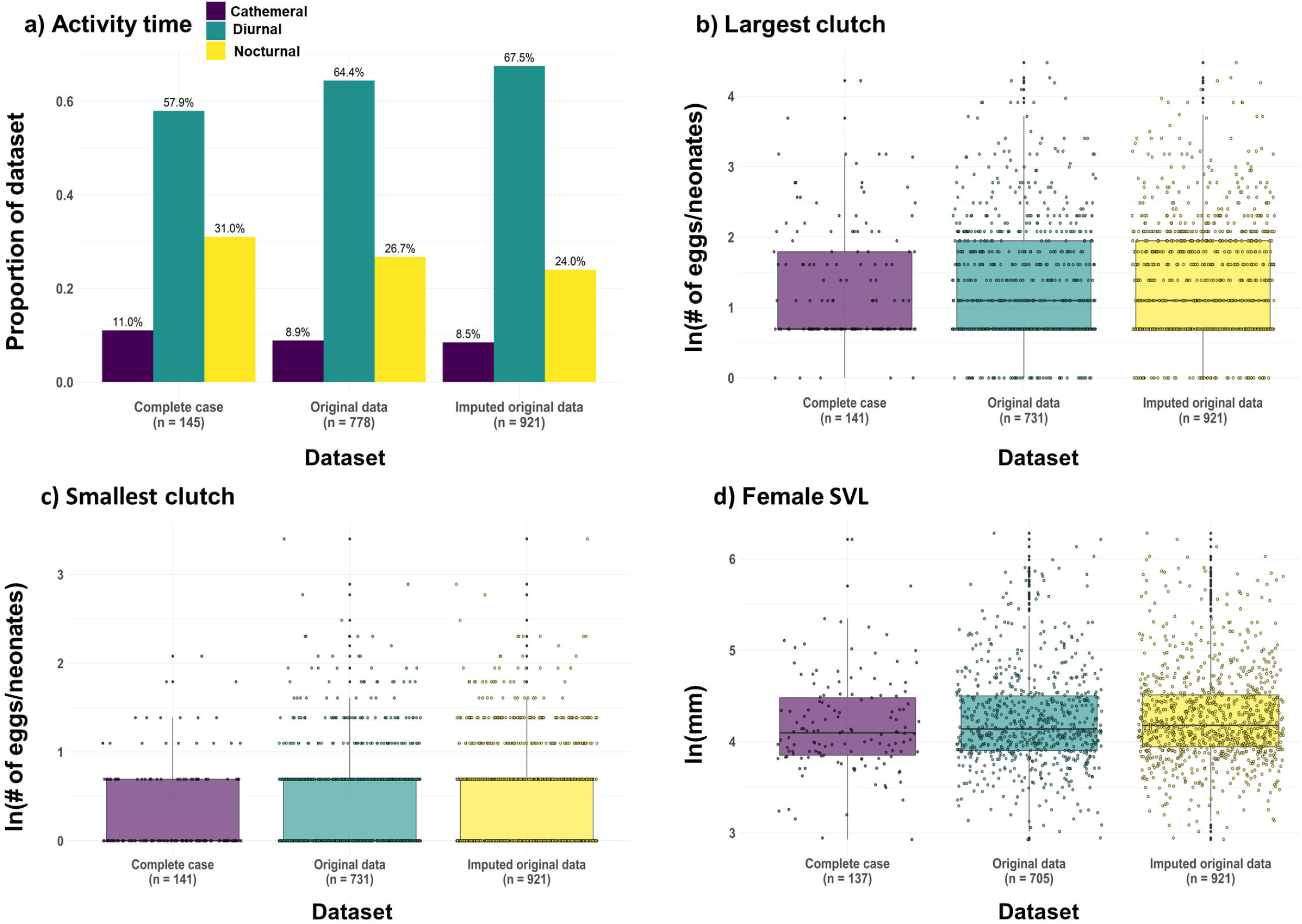
Comparison of quantitative characteristics across datasets. Comparison of a) categorical frequencies for the trait activity time and distributions for the traits b) largest clutch, c) smallest clutch, and d) female SVL of the complete-case data, original data, and imputed data. The natural logarithm (ln) of the numerical data were taken to improve visualization.

## Discussion

To select a best-suited imputation method for the given squamate trait dataset, performances for four candidate methods were evaluated in various scenarios: under different missingness mechanisms and both with and without phylogeny. In alignment with previous evaluations of imputation methods using trait data [11,34,44], the method that performed best (resulted in the lowest error rates) varied in accordance with the simulation settings. When the candidate methods are considered without phylogeny (Fig. 2), as may be the case when molecular data for a taxon are unavailable or of low quality, the best overall method was *MICE. MICE* demonstrated strong performances in previous evaluations of imputation techniques in mammalian [11] and plant [44] trait datasets. The robustness of the predictive mean matching model is appealing for the non-linear relationships and non-normal distributions commonly observed in numerical trait data [45,46]. This may explain the superior performance of *MICE* in the case of smallest clutch, a count trait with a right-skewed distribution (many species with smallest clutch size = 1). Predictive mean matching has also been shown to perform well on smaller sample sizes [46], as seen in the current study (*n* = 152). Its use in trait imputation is therefore an appealing option when sequence data are scarce or phylogenetic signal in the data is weak.

As reported in previous studies [11], imputation error rates tended to increase with missingness proportion and varied amongst different traits. Indeed, for some traits at high missingness percentages, methods performed similarly or worse than mode/mean imputation. These poor performances may be explained by the distributional properties of the trait in question; for example, when considering the trait activity time (see Fig. 2a), a loss of information in this multi-categorical trait due to large proportions of missing data (30-40%) reduced the imputation accuracy for *RF*. This is an issue that can be mitigated through the addition of phylogeny, as imputation accuracy was improved for activity time at high missing proportions when information from any tree was added (see Fig. 3a). However, adding phylogenetic information does not always improve imputation performance; on the contrary, in some instances its inclusion led to increased error rates. The effect of phylogeny therefore appears to be situational and linked to the method used, the underlying mechanism of the missingness in the data, and quantitative attributes and evolutionary history of the target trait.

As phylogenetic resolution varies between nuclear and mitochondrial gene trees, the number of eigenvectors used for imputation varied in accordance. In this study, the threshold method was used to determine the number of eigenvectors to be included; however, it is possible that the use of too many eigenvectors (i.e. > 40) [47], with less information provided by each eigenvector, can introduce more noise or lead to overfitting. *MICE* was particularly sensitive to changes in the number of eigenvectors, as the inclusion of many (> 20) tended to drastically increase error rates. A similar increase in error rate was found in Johnson *et al*. [34] when phylogenetic information was added to *MICE*, especially under MNAR (e.g. larger values more likely to be missing). Penone *et al*. [11] restricted their maximum number of eigenvectors to 10 and suggest that the use of too many eigenvectors can mask the information provided by other traits in the imputation process. Indeed, in the current study, *MICE* performed well when the number of predictors were low, as in the case of trait-only imputation and when the number of eigenvectors were limited. Thus, analyses using phylogenetic eigenvectors for imputation may consider the use of tree-based methods such as *RF* (or recursive partitioning; see Kim *et al*. [32]) that are more robust to high-dimensional data.

The best-suited method overall for the given dataset was identified as *RF* with c-mos. This method resulted in the lowest error rates for the majority of traits, when all missingness mechanisms were considered. This result supports previous evaluations of the effectiveness of *RF* for mixed-type data [25]. For all methods, adding phylogenetic information reduced imputation error rate for traits of all types (binary, categorical, count, continuous); however, the amount of error reduction varied according to the method and tree used. For example, trees built using nuclear-derived phylogenetic information (i.e. c-mos, RAG1, MG) generally conferred a greater improvement in imputation performance relative to mitochondrial COI for the trait insular endemic; on the contrary, COI information was more useful for the trait smallest clutch. Due to their faster rates of nucleotide substitution, mitochondrial genes are less adept at resolving deeper phylogenetic relationships relative to nuclear genes [48]. However, COI often still conferred a reduction in imputation error rate, in some cases more so than the nuclear genes; mitochondrial sequences therefore should be used when nuclear data are unavailable and may be advantageous when studying more closely related species. Overall, these results demonstrate the applicability of single gene trees for imputation when high-quality sequence data are scarce, as may be the case for understudied taxa.

Strength of phylogenetic signal also appeared to correlate with error ratio (i.e. the magnitude of performance enhancement) for categorical traits. The same pattern was not as apparent for numerical traits, however. This may stem from the limited range of phylogenetic signal observed for these other types: all genes displayed significant levels of phylogenetic signal for all traits, many of which verged toward λ = 1 (higher trait conservation). This may suggest that the boost in performance due to phylogeny is negligible beyond a certain level of phylogenetic signal. However, imputation of a greater number and variety of traits that do not display any evidence of phylogenetic signal would need to be included to test this assertion.

The comparison between the distributions and categorical frequencies of the complete-case, original, and imputed trait data support the efficacy of imputation for mixed-type data when the method is carefully chosen. A greater than 6-fold increase in sample size when using imputed data (*n* = 152 for complete-case vs. *n* = 921 for imputed data) is striking and illustrates the information loss that can occur when using a complete-case approach. Complete-case data often do not capture the true variability of the data; instead, they comprise a biased subset and, in turn, the potential for erroneous inferences. Previous studies using clinical data [49] and mammalian trait data [11] found that inferences derived from imputed datasets are less biased when compared to those obtained using complete-case datasets. However, the missing values in these studies were introduced either completely at random (MCAR) or at random (MAR). Although imputation performs well under MCAR and MAR, the mechanism of missingness is often difficult to determine in practice [23,50]. Imputation has been shown to perform poorly in scenarios with biased missingness, such as when extreme values or values in the tails of the distribution of the population are disproportionately missing [34]. If data are truly MNAR and the imputation method is not carefully chosen, imputed values and the inferences derived therein may be inaccurate. A recent study completed by Jardim *et al*. [51] suggests that accurate estimation of phylogenetic signal from imputed datasets is contingent on several variables, including the amount of missing data, missing mechanism, and the evolutionary trajectory of the trait itself. For example, in the current study, as values closer to the equator were missing for the latitude trait under MNAR, mean imputation outperformed most other imputation methods. Due to the prevalence of allopatric speciation modes in diversification [52,53], closely related species can inhabit different latitudes or distributions; traits with such evolutionary histories may be less suitable for imputation. Therefore, we agree with Johnson *et al*. [34] and Jardim *et al*. [51] that caution should be taken and the properties of the dataset of interest be inspected prior to and following imputation. Testing imputation methods using a real data-driven simulation strategy as we demonstrate here would provide useful insight as to whether imputation is a suitable alternative to complete-case analysis.

As is often the case when constructing a complete-case dataset, several traits were excluded from this study. These included many categorical traits that were invariant in the complete-case dataset, such as those containing information about geography or habitat. In turn, the range of phylogenetic signal for traits was also limited. It was therefore not feasible to truly gauge the relationship between error ratio and phylogenetic signal strength in traits as they all exhibited significant levels. The continued collection of high-quality trait data for both known and novel species is necessary to further probe these types of relationships. For instance, in the case of Squamata, snake species are disproportionally undersampled [9] and were thus not included in the current study. An increase in data availability would also facilitate additional research on the use of imputation methods in real datasets. Simulated trait data do not fully capture the nuances of real datasets, and comparative evaluations using real data and different taxonomic groups are needed to test whether imputing values is practical, particularly in cases of biased missingness.

Missingness in datasets is a pervasive issue in the realm of biological research. It is particularly problematic for those taxonomic groups threatened by extinction, or that are small or reside in understudied areas of the globe. As trait data can take on many forms, methods that can accurately predict missing values for diverse data types are instrumental for the study of these obscure groups. Previous work has focused largely on numerical data, and consideration of imputation performance for categorical traits is necessary to drive research in this field forward. Every real trait dataset comprises a unique data structure shaped by both ancient and contemporary processes, including the evolutionary history of the traits and existing sampling biases. Consequently, some datasets may be more suitable for a particular imputation method than others, especially so in the case of phylogenetic imputation. Researchers should therefore take care to understand the properties of their dataset prior to and following imputation. Here, we use a real data-driven selection strategy to select a best-suited imputation method for a given target dataset. Simulating missingness using real data more accurately reflects the characteristics and nature of the original data. Furthermore, our results suggest that, given a carefully selected suitable imputation method, imputation can be used to bolster sample size while simultaneously preserving the distributional properties of the original data. Derived inferences may then more accurately represent the biological phenomena under investigation.

## Materials and methods

### Trait data

Traits are defined here as characteristics that are typical of a species. These may refer to characteristics relating to the biology of a species or the environment in which it resides. Data for squamates (lizards and amphisbaenians; order: Squamata) were selected for analysis as complete-case observations were available for at least 100 species as well as both categorical and numerical traits. In addition, these species had DNA sequence records publicly available for both mitochondrial and nuclear markers. Squamata represent an incredibly diverse group of vertebrates (∼10,000 species) [54], inhabiting disparate environments and boasting a broad range of morphological features. However, trait data for Squamata are undersampled relative to mammal and bird groups, particularly in tropical regions that are home to diverse species at risk [9]. As of 2022, 19.6% of squamate species are estimated to be under threat of extinction [55]. Imputation may offer additional avenues to identify those traits correlated with risk status in squamates [56,57] and in doing so, contribute to biodiversity conservation efforts in vulnerable areas. Trait data were obtained from a dataset published by Meiri [16,17] (other datasets were also considered, see S1 File). This dataset contains information about the habitat, life history, morphology, behaviour, and conservation threat level of 6,657 squamate species (lizards and amphisbaenians, not including snakes) [16,17]. The raw trait data were downloaded into R v. 4.0.3 [58].

To assess which types of genetic information are useful for imputation, sequence data for both mitochondrial and nuclear markers were used. The Barcode of Life Data System (BOLD) [59] was used as the source for mitochondrial sequence data as it contains thousands of published cytochrome *c* oxidase subunit I (COI) partial gene sequence records (16,676 sequences for over 2000 Squamata species as of July 16^th^, 2021). COI barcode sequence data [60] for Squamata were obtained from a published dataset [61], originally downloaded into R on March 12^th^, 2020. Data were filtered for records that have been identified to the species level, as this information was necessary for trait matching purposes. Additional quality control checks on the sequence data included trimming N and gap content from sequence ends and removing sequences with greater than 1% of internal N and/or gap content across their entire sequence length. Sequences between 650 and 1000 bp were retained to facilitate downstream multiple sequence alignment. As multiple COI sequence records are available for many species, a centroid sequence selection process was employed to find a typical representative sequence for each species [62]. The centroid sequences used for multiple sequence alignment have been made available at [63]. The *AlignTranslation* function from the R package “DECIPHER” v. 2.18.1 [64,65] was used to perform a multiple sequence alignment on the centroid sequences. *AlignTranslation* was used as it performs a multiple sequence alignment guided by the translated amino acid sequence, which is more reliable than an alignment based on nucleotide data alone [64]. The translated final alignment was visualized using the *ggmsa* function from the R package “ggmsa” v. 0.06 [66] to verify the nucleotides were in the correct reading frame and to check for the presence of stop codons. The COI multiple sequence alignment and corresponding process IDs are available on Dryad [67]. Nuclear sequence data were obtained from a multigene alignment published in Pyron *et al*. [68,69]. This alignment is comprised of sequence data for 12 genes (seven nuclear, five mitochondrial) and 4161 species of Squamata [68]. The alignment was partitioned into its constituent gene alignments using RAxML v. 8 [70].

To form a complete-case dataset for downstream missingness simulations, species names from the COI alignment were first matched against the species names in the trait dataset. Those species that had available data for at least five traits (both categorical and numerical) and a corresponding COI sequence record were then matched against the species names in the nuclear multigene alignment. The nuclear markers oocyte maturation factor (c-mos) and recombination activating gene 1 (RAG1) had the largest number of available records for the species in the complete-case dataset and were selected for analyses. Corresponding sequence identifiers of those records selected (either BOLD process ID [59] or GenBank accession number [71]) available in S1 Table and on Dryad [67]. Final checks were performed on the trait data in the complete-case subset. Categorical traits with severe class imbalances and very low variability (e.g. more than 90% of observations in one of the categories and/or the remaining observations sparsely dispersed across other categories), such as reproductive mode, geographic range, and substrate, were excluded from the study. The distributions of numerical trait data were visualized to check for the presence of severe outliers. For each numerical trait, an upper threshold was calculated as follows: quartile 3 + (3 × the interquartile range of the data). Severe outliers are defined here as those values that exceed the upper threshold. If identified, these values were verified in the primary literature to ensure they were real datapoints and not the result of data entry error. The final dataset contains information for the seven most complete traits, including the multi-categorical trait activity time, the binary trait insular endemic, the count traits: largest clutch and smallest clutch, and the continuous traits female snout-vent length (SVL), maximum snout-vent length (SVL), and latitude (geographic centroid for the species [72]). The final dataset comprised 152 species, representing 25 Squamata families (S2 Table). To maintain a sufficient sample size for missingness simulations, it was necessary to reduce the stringency for inclusion of species in the complete-case dataset; otherwise, the sample size will drop to 121 if only species without missing values in their traits are included. Consequently, we permitted some missing values (no more than 10% for each trait) present in the so-called “near complete-case dataset”. To ensure enough data were available for each species for imputation, missing values were only permitted in traits for species that had observed values for at least three of the other traits in the dataset. For further details on these traits, see S3 Table.

### Phylogenetic information

As high-quality sequence data may be unavailable for multiple markers for understudied taxa, one of our objectives was to demonstrate the utility of single gene trees for imputation. The alignments for the COI, c-mos, and RAG1 sequences were used to build maximum likelihood gene trees in RAxML v. 8 [70]. For comparison purposes, a composite gene tree was also built using a combined multigene alignment of the COI, c-mos, and RAG1 sequences (referred to as “MG” for multigene). The model GTRGAMMAI was specified (option -m), and the alignments were partitioned based on codon position (option -q). The trees were then read into R and made ultrametric using the *chronos* function in the R package “ape” v. 5.4.1 [73]. Phylogenetic eigenvectors were extracted from each tree and for each trait using the “MPSEM” package v. 0.3.6 in R [74]. The number of eigenvectors that explained greater than or equal to 65% of the phylogenetic structure variance was used (see S1 File for further details on this process). Following the method of Penone *et al*. [11], the phylogenetic eigenvectors were appended to the near complete-case dataset and treated as predictors in the model to impute the missing value of a given trait. All trees used in the phylogenetic imputation process have been made available on Dryad [67].

Previous studies have suggested that phylogenetic signal strength in simulated trait data is positively correlated with imputation accuracy [32,75]. To assess this association using real data, we measured phylogenetic signal for each trait using Pagel’s λ [42] for numerical traits and the *D* metric [41] for categorical traits. Pagel’s λ is estimated using maximum likelihood and represents the value that optimally transforms a phylogenetic variance-covariance matrix to fit the observed trait data structure. A λ value of 0 indicates no phylogenetic signal (star-shaped phylogeny), whereas a λ value of 1 suggests that the trait data adhere to a Brownian motion (BM) model of evolution [42]. The *D* metric represents whether the number of transitions of a binary trait varies from the expected number under a BM model [41]. A *D* value of 0 indicates that the trait data adhere to a BM model, and a *D* value of 1 indicates that there is no phylogenetic signal in the trait data. A *D* value greater than 1 signifies phylogenetic overdispersion. Alternatively, a *D* value less than 0 suggests the trait is phylogenetically conserved [41]. These metrics were calculated separately for each trait using each gene tree (S1 File). The *phylosig* function in the R package “phytools” v. 0.7.70 [76] and *phylo*.*d* function in the R package “caper” v. 1.0.1 [77] were used to measure λ and *D*, respectively.

### Candidate imputation methods

The performances of four candidate imputation methods were considered: mean/mode imputation, *k*-nearest neighbour (*KNN*) (“VIM” package v. 6.1.0) [24], random forests (*RF*) (“missForest” package v. 1.4) [40,78], and multivariate imputations by chained equations (*MICE*) (“mice” package v. 3.13.0) [27]. Mean (for numerical traits) / mode (for categorical traits) imputation, the simplest method, was used as a baseline for comparison. The remaining methods were chosen due to their popularity in trait-based studies [35,79] and capacity to impute both numerical and categorical traits. These methods have also been evaluated in previous studies of trait data imputation [11,34,44]. *KNN* and *RF* are single imputation methods as they provide a single estimation of the missing value. *MICE* is a multiple imputation method that performs imputation *m* times on the dataset with missing values, resulting in *m* imputed datasets. The *MICE* algorithm utilizes chained equations to estimate missing values and offers several different models for imputing data. In this study, the predictive mean matching model was used to estimate missing continuous data. Predictive mean matching is the default model for continuous data in *MICE* and performed well in previous evaluations using trait data [44,80]. Predictive mean matching fills the missing observation with a random value selected from a “donor” pool for the missing observations. This pool is created by fitting a regression model on the observed data and selecting *k* fitted values that are closest to the predicted value for the missing observation [45,46]. Logistic regression is a common approach for predicting missing binary data and is the default method for data of this type in *MICE*. Logistic regression and polytomous logistic regression models were thus used to impute values for the binary trait insular endemic and the nominal multi-categorical trait activity time, respectively. To obtain a final imputed value for *MICE*, the mean and mode values were taken across the *m* datasets for numerical traits and categorical traits, respectively. See S1 File for further details on imputation algorithms.

Missing values of each trait (“target trait”) are imputed using values of the other traits (“auxiliary traits”); however, not all of the auxiliary traits are useful for imputation. Association tests between each pair of traits were used to filter out irrelevant auxiliary traits and build a more parsimonious imputation model for the target trait. Regression models were used in the association tests in which the target trait was specified as the response variable and each one of the auxiliary traits was specified as the covariate. Linear regression, Poisson regression, logistic regression, and polytomous logistic regression models were used for continuous, count, binary, and multi-categorical target traits, respectively. Only auxiliary traits with a coefficient not significantly equal to zero were retained in the imputation model for a particular target trait. Finally, as methods such as *KNN* are sensitive to the range of the data, numerical traits were natural log-transformed prior to imputation.

### Real data-driven simulation strategy

#### Missingness simulations

The first step in the real data-driven simulation strategy was to simulate missing values in the near complete-case dataset (Fig 7). Three different missingness mechanisms were considered: 1) missing completely at random (“MCAR”); 2) missing at random (“MAR”); and 3) missing not at random (“MNAR”). Under the MCAR mechanism, missing values were randomly introduced into the near complete-case dataset at different proportions (0.10, 0.20, 0.30, and 0.40). In cases where traits had values that were already missing (up to 10%), missing values were introduced on top of these (i.e. up to 50% missingness). To reduce stochasticity and maintain a fair comparison of imputation performance across different missing proportions, and not introducing variability relating to species identity, missing data for each increase in proportion (e.g. from 0.10 to 0.20 missingness) were added upon the missing values of the previous proportion. To simulate MAR using real data, logistic regression models were fitted to the original Meiri [16] dataset (*n* = 6657) to identify which auxiliary traits were significantly associated with the missingness for each target trait. In the fitted model, the indicator of whether an observation is missing or not was treated as the response variable and auxiliary traits specified as predictors. The fitted models were then used to introduce missing values into the near complete-case dataset (for further details see S1 File). To test how the imputation methods perform in cases of extreme bias, MNAR scenarios were simulated for each trait. Values were removed from the 10^th^ percentile of the tail of data distribution for numerical biological traits, e.g., the 10^th^ percentile of the lower latitudes (between 10° and -10°); and from a single category for categorical traits, e.g., “nocturnal” category for activity time and “yes” category for insular endemic. These values were removed to emulate realistic MNAR scenarios for Squamata (see S1 File for further information).

**Fig 7.**
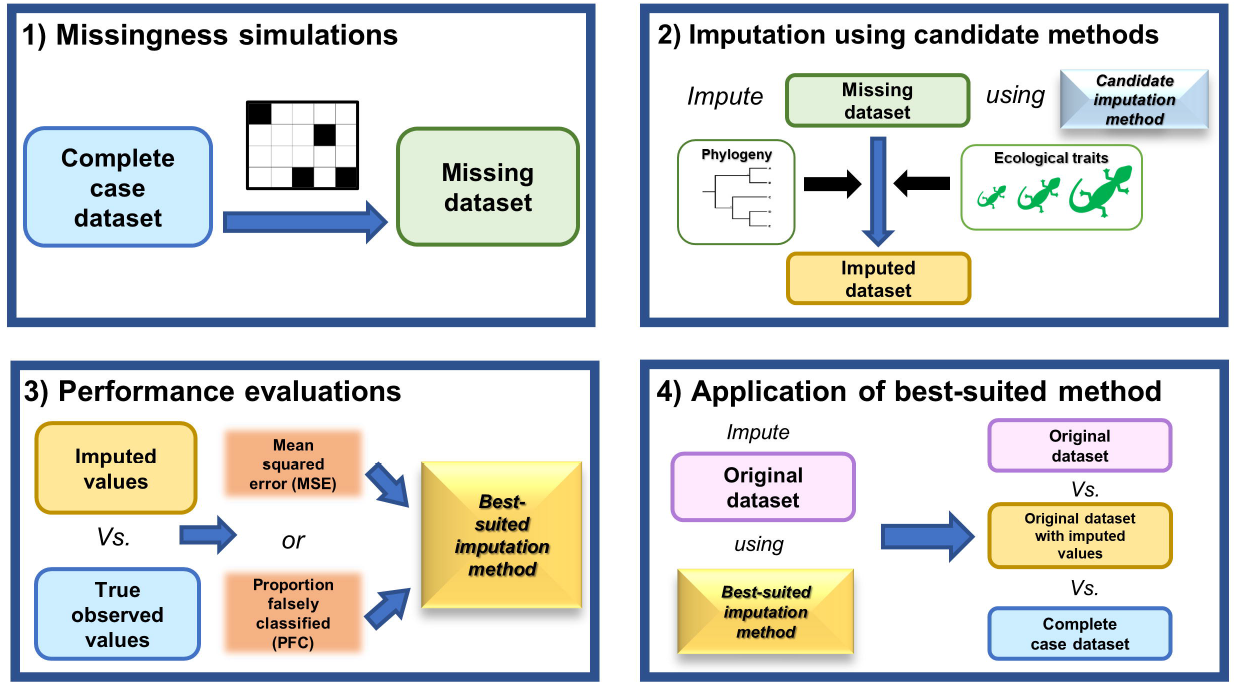
Workflow of the real data-driven simulation strategy. 1) Missing values are simulated in the near complete-case dataset under MCAR, MAR, and MNAR mechanisms. Missing dataset refers to the near complete-case dataset with simulated missing values; 2) imputation of simulated missing values using different candidate methods (with or without phylogeny); 3) identification of the best-suited method based on the performances (MSE and PFC) of the candidate methods. The imputed values from 2) are compared to the true observed values in the near complete-case dataset to derive MSE and PFC for numerical and categorical traits, respectively. Note that the “true observed values” refer to the trait values reported in the source Meiri (2018) dataset [16,17]. These values are assumed to contain a negligible amount of measurement error in this context and provide a close representation of the true biological values for squamates. MSE and PFC are averaged across 100 replicates; and 4) application of the best-suited imputation method to the original target dataset with missing values. Original data, original data with imputed values, and complete-case data characteristics and distributions are compared to assess how imputation impacts dataset properties.

#### Imputation using candidate methods

The second main step in the strategy was to apply each candidate method to impute the simulated missing values under the different simulation settings. A range of parameters and their values were considered for the different candidate imputation methods (see S1 File for details on this process). The parameters that resulted in the lowest error rate were used in the imputation model. Imputations using only trait data were first performed on the simulated missing dataset. Imputations were again performed using trait data and phylogenetic eigenvectors derived from either COI, RAG1, c-mos, or MG trees. This amounted to 96 different combination settings with respect to method and missingness mechanism. The entire process was repeated 100 times for each combination of settings, resulting in 9,600 runs of the simulation and imputation pipeline strategy.

#### Performance evaluations

The third main step in the strategy was to evaluate the performances of each candidate method under the different simulation settings. Imputation performance is defined in this context as how well (i.e. accurately) the candidate method predicts the true observed values. The “true observed values” refer to the values as reported in the Meiri (2018) dataset [16,17]. It is assumed that these values are close to the true biological values for squamates and that the error in the reported values is generally small in comparison with the total variability in each trait across all species in the dataset. To assess performance, mean squared error (MSE) rates and proportion falsely classified (PFC) rates were computed for numerical and categorical traits, respectively. To compute MSE and PFC, only those imputed values that were simulated as missing in the near complete-case dataset and that have true observed values were used. For both metrics, values closer to 0 are indicative of better performance. To illustrate the impact of missingness percentage on imputation accuracy, we generated an equal number of missing data replicates for each missingness percentage (10%, 20%, 30%, 40%). As other simulation settings (missingness mechanism, imputation method with or without phylogenetic information) are preserved for each missingness percentage, this enables a fair comparison of imputation accuracy impacted by missingness percentage. Vice versa, the performance of imputation using different methods can be compared fairly using replications of the same missingness percentage. The performance metrics (MSE and PFC) were averaged across the 100 replicates for each combination of methods for each trait. The packages “ggplot2” v. 3.3.5 [43] and “plotly” v. 4.10.0 [81] were used to visualize results in R. The best-suited method is defined here as the imputation method, either with or without phylogeny, that resulted in the lowest average error rates (either MSE for numerical traits or PFC for categorical traits) for the most traits (i.e. “a majority vote”). To select the best-suited method, the results of the MAR simulations were first considered as these mimic realistic biological scenarios. In case of more than one method performing similarly under MAR (e.g. a tied vote), the results of MNAR and MCAR were also considered.

#### Application of best-suited imputation method to original dataset

The fourth main step in the strategy was to apply the best-suited imputation method to the original target dataset and compare the characteristics between the original data, original data with imputed values, and complete-case data for each trait. To assess the influence of imputed values on data characteristics, summary statistics for each trait were calculated using the original dataset with imputed values and qualitatively compared with the summary statistics of both the original and complete-case data. To investigate whether the phylogenetic information improves the imputation accuracy for a given trait and imputation method, the following error ratio was calculated for each trait and each method:

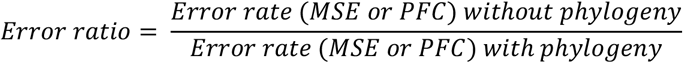

An error ratio value greater than 1 indicates an improvement in imputation performance resulting from the addition of phylogenetic information. To observe the trend of the effect of phylogenetic signal strength on the imputation of different traits, the error ratio values were plotted against the λ and *D* metrics for numerical and categorical traits, respectively.

## Acknowledgements

We would like to thank Dr. Cameron Nugent and Dr. Karl Cottenie for their helpful insights regarding the design and structure of the simulation pipeline. Thank you to Dr. Angela Canovas and Dr. Mark Engstrom for their thoughtful suggestions on the methodology and manuscript. We also thank Dr. Tyler Elliott for his valuable comments on the manuscript and code. Finally, we thank the many researchers who have collected trait and sequence data and made them publicly available. This work would not have been possible without you.

## Supporting information

**S1 File. Supplementary Information**.

**S1 Fig. Phylogenetic signal measurements**. Measures of phylogenetic signal for a) categorical and b) numerical traits in gene trees constructed for mitochondrial COI and nuclear c-mos and RAG1. Asterisks indicate significance at the 0.05 level, according to results from hypothesis tests comparing the results to a null model (no phylogenetic signal). Fritz and Purvis’ *D* metric [41] and Pagel’s λ [42] were used to measure phylogenetic signal for categorical and numerical traits, respectively. In the case of *D*, lower values are indicative of higher levels of phylogenetic conservation for the trait; conversely, higher values of λ suggest stronger phylogenetic signal. As the *D* metric only measures the phylogenetic signal of binary traits, the three-level categorical trait AT was broken down into the binary traits “AT: Diurnal” and “AT: Nocturnal”.

**S1 Table. Sequence identifiers**.

**S2 Table. Taxonomic composition of near complete-case trait dataset (n = 152)**.

**S3 Table. Descriptions and additional details for traits in the near complete-case dataset**.

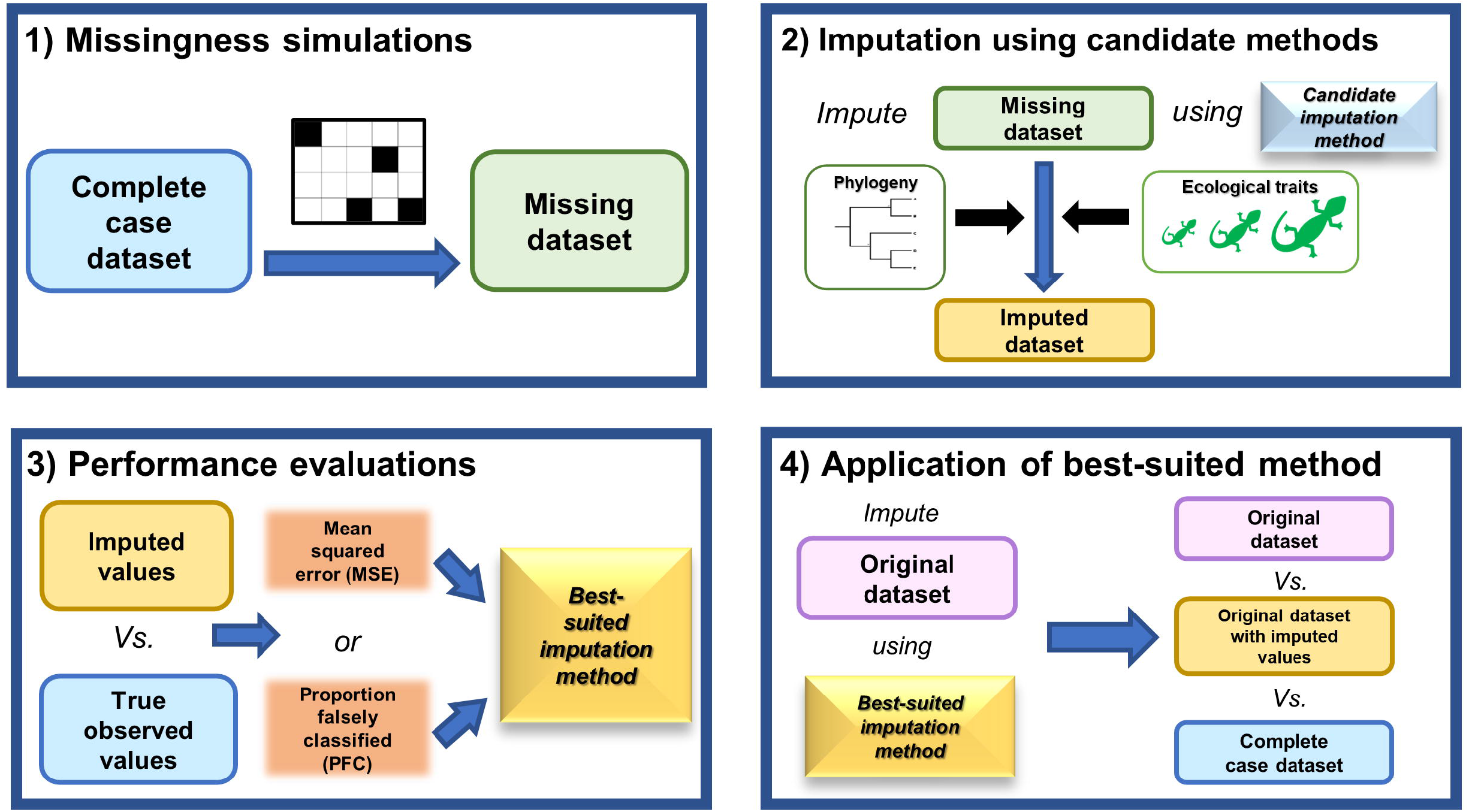

