## Supplementary figures and images for "A real data-driven simulation strategy to select an imputation method for mixed-type trait data"

### S1 Figure

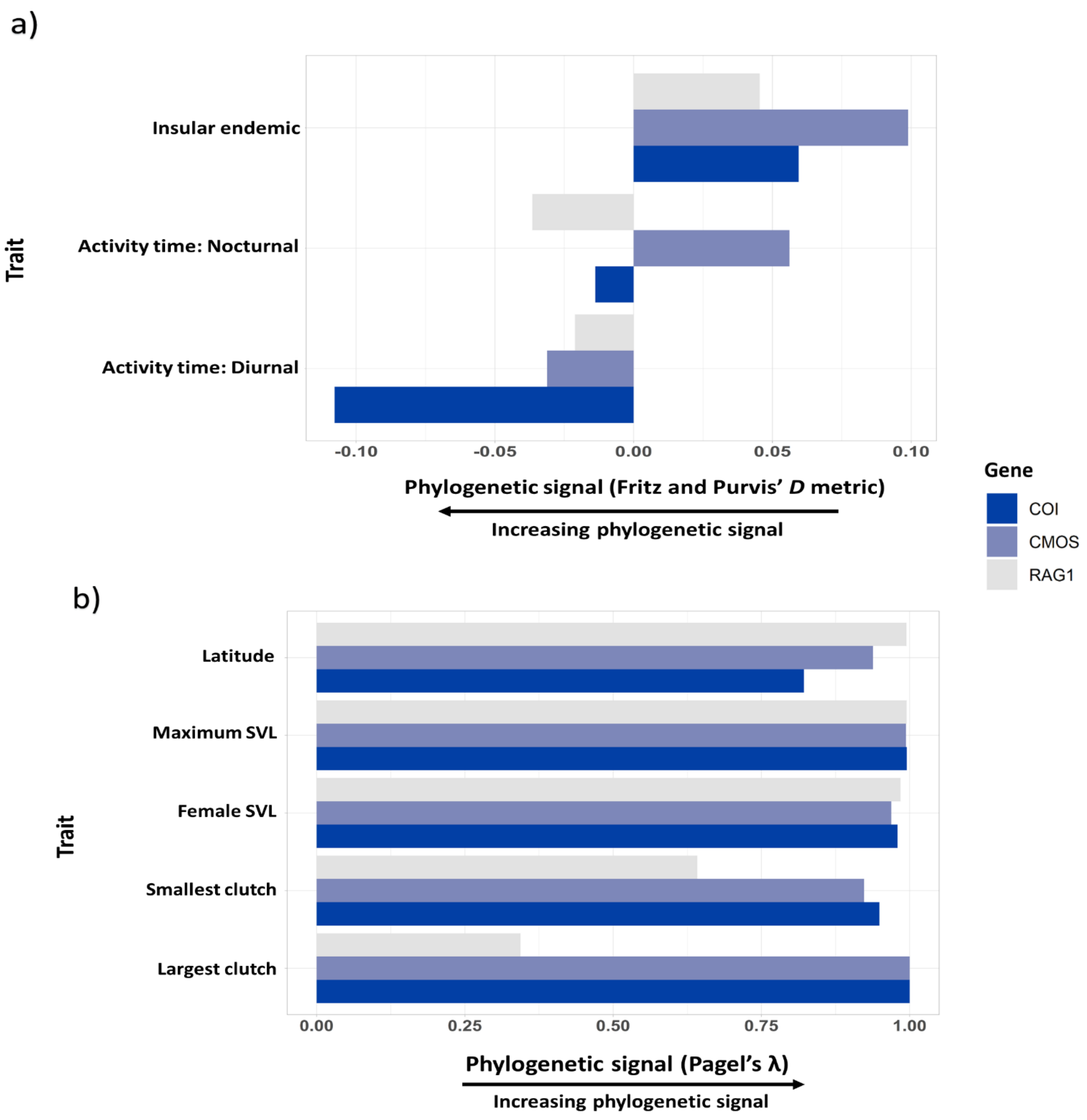
