## Supplementary material for "A real data-driven simulation strategy to select an imputation method for mixed-type trait data": S1 Table

**S1 Table. Sequence identifiers.**

| Species name | COI Process ID | c-mos Accession Number | RAG1 Accession Number |
|---|---|---|---|
| *Acanthodactylus boskianus* | NPLRP211-08 | EF632251 | EF632206 |
| *Aeluroscalabotes felinus* | GBMNA11757-19 | HQ426517 | JN654855 |
| *Agama agama* | BACOR025-13 | AF137530 | EU402825 |
| *Agamura persica* | ABLRP470-07 | DQ852728 | JQ945281 |
| *Alsophylax pipiens* | NPLRP398-08 | JQ945531 | JQ945284 |
| *Anniella pulchra* | EANAH930-12 | AY487350 | AY662605 |
| *Apathya cappadocica* | GBGC12728-13 | EF632268 | EF632223 |
| *Aprasia parapulchella* | GBMTG4679-16 | AY134539 | HQ426260 |
| *Bipes biporus* | GBMTG502-16 | AF039482 | AY662616 |
| *Bipes canaliculatus* | GBMNA11783-19 | FJ518700 | FJ518701 |
| *Blaesodactylus antongilensis* | REPT235-12 | JQ945534 | EU054230 |
| *Blanus cinereus* | GBMTG1541-16 | DQ324864 | EU108523 |
| *Brookesia ambreensis* | REPT049-12 | FJ984313 | FJ984243 |
| *Brookesia antakarana* | REPT050-12 | FJ984312 | FJ984242 |
| *Brookesia brygooi* | REPT153-12 | FJ984306 | FJ984236 |
| *Brookesia decaryi* | GBMNA11833-19 | FJ984308 | FJ984238 |
| *Brookesia ebenaui* | REPT057-12 | FJ984300 | FJ984230 |
| *Brookesia exarmata* | REPT360-12 | FJ984291 | FJ984220 |
| *Brookesia griveaudi* | REPT371-12 | FJ984321 | FJ984251 |
| *Brookesia minima* | REPT396-12 | FJ984280 | FJ984209 |
| *Brookesia peyrierasi* | REPT236-12 | FJ984287 | FJ984216 |
| *Brookesia stumpffi* | REPT051-12 | FJ984318 | FJ984247 |
| *Brookesia superciliaris* | GBGC6845-09 | FJ984303 | FJ984233 |
| *Brookesia tuberculata* | REPT052-12 | FJ984281 | FJ984210 |
| *Bunopus tuberculatus* | NPLRP073-08 | AF148706 | JQ945287 |
| *Caledoniscincus austrocaledonicus* | GBGCR3727-19 | DQ675404 | EU568024 |
| *Calumma gastrotaenia* | REPT251-12 | FJ984263 | FJ984191 |
| *Calumma nasutum* | REPT233-12 | JN030463 | HQ130637 |

| | | | |
|---|---|---|---|
| *Chalarodon madagascariensis* | GBMNA11850-19 | AY987987 | FJ356745 |
| *Chlamydosaurus kingii* | GBMNA18857-19 | DQ340665 | JF806191 |
| *Cnemaspis limi* | GBMNA11758-19 | EF534935 | EF534809 |
| *Coleonyx variegatus* | GBMTG991-16 | EF534901 | EU108526 |
| *Crossobamon orientalis* | ABLRP217-07 | DQ852730 | JQ945299 |
| *Crotaphytus collaris* | EANAH716-12 | AY987985 | FJ356749 |
| *Cyrtodactylus irregularis* | GBGCR6135-19 | JQ945551 | JQ945302 |
| *Cyrtopodion scabrum* | DJIB114-17 | HQ426532 | HQ426275 |
| *Dibamus novaeguineae* | GBGC9701-09 | EF450999 | EU108529 |
| *Dipsosaurus dorsalis* | EANAH605-12 | AF148705 | FJ356747 |
| *Ebenavia inunguis* | BACOR034-13 | FJ830144 | EF536143 |
| *Eremias arguta* | NPLRP542-08 | EF632258 | EF632213 |
| *Eublepharis macularius* | GBGCR6235-19 | EU366458 | EF534776 |
| *Eutropis multifasciata* | IQHLM132-06 | DQ238978 | AY444055 |
| *Furcifer lateralis* | REPT394-12 | FJ984272 | JQ073208 |
| *Furcifer polleni* | BACOR067-13 | FJ984267 | JQ073215 |
| *Furcifer willsii* | REPT250-12 | HQ130551 | HQ130640 |
| *Geckolepis maculata* | BACOR081-13 | JQ945562 | EU054211 |
| *Gehyra mutilata* | NPRPV022-08 | FJ830146 | FJ830237 |
| *Gekko chinensis* | GBMNA11763-19 | JQ945571 | JN019123 |
| *Gekko gecko* | NPRPV009-08 | EU366455 | AY662625 |
| *Gekko vittatus* | GBGC10139-09 | JQ945575 | JN019137 |
| *Gonatodes albogularis* | GBGCR6260-19 | EF564078 | EF534797 |
| *Goniurosaurus luii* | GBMNA11755-19 | HQ426538 | HQ426287 |
| *Heloderma suspectum* | GBGCR449-15 | AY662566 | AY662606 |
| *Hemidactylus flaviviridis* | DJIB124-17 | HQ426541 | HM559694 |
| *Hemidactylus frenatus* | GBGC10802-13 | EF534940 | EF534814 |
| *Hemidactylus mercatorius* | BACOR085-13 | AY863046 | JQ073244 |
| *Hemidactylus platycephalus* | BACOR077-13 | AY863045 | JQ073246 |
| *Hemidactylus platyurus* | NPRPV024-08 | HQ426530 | HM559685 |

| | | | |
|---|---|---|---|
| *Hemidactylus robustus* | DJIB231-17 | HQ426549 | EU054271 |
| *Hemitheconyx caudicinctus* | GBMNA11756-19 | HQ426552 | HQ426294 |
| *Hemitheconyx taylori* | GBGC11800-13 | HQ426553 | HQ426295 |
| *Heteronotia binoei* | GBGC4418-08 | JQ945580 | EU054285 |
| *Homonota fasciata* | GBMNA18955-19 | EU293674 | EU293629 |
| *Lacerta agilis* | FBHER281-14 | EU365405 | EF632222 |
| *Lacerta viridis* | GBMTG853-16 | DQ097131 | EU108535 |
| *Lampropholis guichenoti* | GBGCR3728-19 | DQ675352 | EU568111 |
| *Leiolepis belliana* | GBGCR211-15 | FJ984253 | AY662587 |
| *Lepidodactylus lugubris* | GBMNA11768-19 | EF534938 | EF534812 |
| *Lepidophyma flavimaculatum* | GBMTG982-16 | EU116715 | EU108567 |
| *Lygodactylus miops* | REPT023-12 | HQ426556 | HQ426299 |
| *Lygodactylus mirabilis* | REPT385-12 | HQ426557 | HQ426300 |
| *Madascincus melanopleura* | REPT223-12 | AY802768 | HM161147 |
| *Marmorosphax tricolor* | GBGCR3729-19 | DQ675367 | EU568023 |
| *Matoatoa brevipes* | REPT295-12 | JQ945587 | EF490724 |
| *Mesalina guttulata* | NPLRP518-08 | EF632274 | EF632231 |
| *Morethia adelaidensis* | GBGCR3730-19 | DQ675368 | EU568109 |
| *Nannoscincus mariei* | GBGCR3731-19 | DQ675372 | EU568021 |
| *Oligosoma microlepis* | GBGCR3874-19 | DQ675375 | EU568088 |
| *Oligosoma smithi* | GBGCR3978-19 | DQ675386 | EU568090 |
| *Oligosoma suteri* | GBGCR3996-19 | DQ675387 | EU568106 |
| *Oligosoma zelandicum* | GBGCR4026-19 | AY818781 | EU568082 |
| *Oplurus cuvieri* | GBGCR1711-18 | EU099677 | AY662601 |
| *Oplurus cyclurus* | REPT095-12 | EU099680 | GU457973 |
| *Paragehyra gabriellae* | REPT083-12 | JQ945603 | JQ945328 |
| *Paroedura androyensis* | REPT383-12 | HQ256721 | EF490721 |
| *Paroedura bastardi* | REPT292-12 | HQ256725 | EF536163 |
| *Paroedura gracilis* | GBGCR282-15 | HQ256726 | EF536161 |
| *Paroedura lohatsara* | GBGCR439-15 | HQ256731 | EF536155 |

| | | | |
|---|---|---|---|
| *Paroedura masobe* | GBGCR440-15 | HQ426560 | EF536145 |
| *Paroedura oviceps* | GBGCR441-15 | HQ256733 | EF536160 |
| *Paroedura picta* | GBMNA11769-19 | EU293692 | EF536150 |
| *Paroedura stumpffi* | GBGCR6441-19 | HQ256740 | EF536154 |
| *Phelsuma abbotti* | REPT364-12 | AY221346 | FJ830148 |
| *Phelsuma antanosy* | REPT097-12 | FJ830064 | FJ830156 |
| *Phelsuma barbouri* | REPT384-12 | FJ830069 | FJ830161 |
| *Phelsuma berghofi* | REPT320-12 | FJ830070 | FJ830162 |
| *Phelsuma breviceps* | REPT003-12 | FJ830072 | FJ830164 |
| *Phelsuma dubia* | BACOR114-13 | FJ830080 | FJ830172 |
| *Phelsuma guimbeaui* | GBMNA18868-19 | AY221372 | FJ830221 |
| *Phelsuma guttata* | REPT020-12 | FJ830085 | FJ830177 |
| *Phelsuma hielscheri* | REPT121-12 | FJ830086 | FJ830178 |
| *Phelsuma laticauda* | BACOR117-13 | AY221343 | FJ830189 |
| *Phelsuma lineata* | REPT001-12 | AY221342 | JN654861 |
| *Phelsuma madagascariensis* | REPT298-12 | EF534937 | EF534811 |
| *Phelsuma malamakibo* | REPT348-12 | FJ830108 | FJ830200 |
| *Phelsuma modesta* | REPT285-12 | FJ830110 | FJ830202 |
| *Phelsuma mutabilis* | REPT352-12 | FJ830114 | FJ830206 |
| *Phelsuma pusilla* | REPT305-12 | FJ830122 | FJ830214 |
| *Phelsuma quadriocellata* | REPT199-12 | FJ830123 | FJ830215 |
| *Phelsuma ravenala* | REPT319-12 | FJ830078 | FJ830170 |
| *Phelsuma serraticauda* | REPT324-12 | FJ830131 | FJ830223 |
| *Phelsuma standingi* | REPT289-12 | FJ830133 | FJ830225 |
| *Phoenicolacerta kulzeri* | GBMNA11795-19 | GQ142151 | GQ142161 |
| *Phrynocephalus mystaceus* | GBGCR6007-19 | AF137527 | GQ242268 |
| *Phyllodactylus unctus* | GBMTG2995-16 | FJ662503 | HQ426312 |
| *Plestiodon skiltonianus* | EANAH819-12 | AF315396 | AY662633 |
| *Plica plica* | GBGCR5143-19 | EF615737 | FJ356742 |
| *Podarcis muralis* | FBHER036-09 | EF632282 | EF632239 |

| | | | |
|---|---|---|---|
| *Pogona vitticeps* | GBMNA11825-19 | DQ340691 | JF806200 |
| *Polychrus marmoratus* | GBMNA11856-19 | AY987983 | FJ356748 |
| *Ptyodactylus guttatus* | GBMNA11775-19 | EU293681 | EU293636 |
| *Ptyodactylus hasselquistii* | NPLRP357-08 | EU293682 | EU293637 |
| *Quedenfeldtia moerens* | GBGCR5605-19 | HQ426574 | HQ426320 |
| *Quedenfeldtia trachyblepharus* | GBGCR5607-19 | EF534930 | EF534804 |
| *Rhineura floridana* | GBMTG497-16 | AY444022 | AY662618 |
| *Saurodactylus mauritanicus* | NPLRP424-08 | EU014324 | EU014356 |
| *Sauromalus ater* | EANAH588-12 | AF315400 | AY662591 |
| *Scincella lateralis* | EANAH892-12 | AY217857 | HM161236 |
| *Scincus scincus* | NPLRP289-08 | AY217873 | HM161238 |
| *Shinisaurus crocodilurus* | GBMNA11809-19 | AY099976 | AY662610 |
| *Stenodactylus sthenodactylus* | ABLRP285-07 | JQ945617 | JQ945339 |
| *Takydromus amurensis* | GBGC11979-13 | EF632287 | EF632244 |
| *Takydromus sexlineatus* | GBMNA11799-19 | EF632288 | EF632245 |
| *Tarentola annularis* | DJIB068-17 | AF363552 | DQ275456 |
| *Tarentola mauritanica* | GBGC12027-13 | AF363566 | EU293641 |
| *Teratoscincus microlepis* | GBGCR6026-19 | EF534926 | EF534800 |
| *Teratoscincus przewalskii* | GBGCR6022-19 | AY662569 | AY662624 |
| *Teratoscincus roborowskii* | ZISPG080-09 | EF534925 | EF534799 |
| *Teratoscincus scincus* | GBGCR6037-19 | EF534927 | EF534801 |
| *Tracheloptychus madagascariensis* | REPT345-12 | DQ100104 | JQ073186 |
| *Tropidurus hispidus* | GBGCR1725-18 | AY987984 | AY988013 |
| *Tropidurus insulanus* | GBGCR5141-19 | EF615738 | EF616390 |
| *Tropidurus oreadicus* | GBGCR1726-18 | EF615739 | EF616391 |
| *Tropiocolotes tripolitanus* | GBMNA11771-19 | JQ945623 | JQ945343 |
| *Uroplatus ebenaui* | GBMNA11773-19 | JN038097 | EF490736 |
| *Uroplatus giganteus* | REPT053-12 | JQ945625 | EF490738 |
| *Uroplatus guentheri* | REPT279-12 | JQ945626 | EF490725 |

| | | | |
|---|---|---|---|
| *Uta stansburiana* | GBMNA11855-19 | AF315389 | DQ385422 |
| *Varanus salvator* | GBGCR4034-19 | AF435017 | EU402828 |
| *Xantusia henshawi* | EANAH907-12 | EU116794 | EU108645 |
| *Zonosaurus madagascariensis* | REPT017-12 | DQ100129 | JQ073185 |
| *Zootoca vivipara* | FBHER093-09 | EF632292 | EF632249 |

Database identifiers for all sequence records used in this study (n = 152 species). Process IDs are included for cytochrome c oxidase subunit I (COI) sequence records obtained from the Barcode of Life Data (BOLD) System (1). GenBank (2,3) accession numbers are included for nuclear oocyte maturation factor (c-mos) and recombination activating gene 1 (RAG1). c-mos and RAG1 sequences were obtained via the multigene alignment published in Pyron *et al.* (4,5).
