## Supplementary material for "A real data-driven simulation strategy to select an imputation method for mixed-type trait data": S2 Table

**S2 Table. Taxonomic composition of near complete-case trait dataset (*n* = 152).**

| Family | Common name | Count (N) | Percentage of dataset (%) |
|---|---|---|---|
| Gekkonidae | Common geckos | 59 | 38.8 |
| Chamaeleonidae | Chameleons | 17 | 11.2 |
| Scincidae | Skinks | 14 | 9.2 |
| Lacertidae | Old world runners/Lacertid lizards | 11 | 7.2 |
| Sphaerodactylidae | N/A | 8 | 5.3 |
| Eublepharidae | Eublepharid geckos | 6 | 3.9 |
| Phyllodactylidae | N/A | 6 | 3.9 |
| Agamidae | Old world arboreal lizards/Agamid lizards | 5 | 3.3 |
| Tropiduridae | Tropidurid lizards | 4 | 2.6 |
| Opluridae | N/A | 3 | 2.0 |
| Bipedidae | Two-legged worm lizards | 2 | 1.3 |
| Gerrhosauridae | Plated lizards | 2 | 1.3 |
| Iguanidae | Iguanas | 2 | 1.3 |
| Xantusiidae | Night lizards | 2 | 1.3 |
| Anniellidae | American legless lizards | 1 | 0.7 |
| Blanidae | N/A | 1 | 0.7 |
| Crotaphytidae | Collared lizards/leopard lizards | 1 | 0.7 |
| Dibamidae | N/A | 1 | 0.7 |
| Helodermatidae | Gila monsters | 1 | 0.7 |
| Phrynosomatidae | North American spiny lizards | 1 | 0.7 |
| Polychrotidae | Anoloid lizards | 1 | 0.7 |
| Pygopodidae | Flap-footed lizards | 1 | 0.7 |
| Rhineuridae | North American worm lizards | 1 | 0.7 |
| Shinisauridae | N/A | 1 | 0.7 |
| Varanidae | Monitor lizards | 1 | 0.7 |

Trait data obtained from Meiri (2018, 2019).
