## Supplementary material for "A real data-driven simulation strategy to select an imputation method for mixed-type trait data": S3 Table

**S3 Table. Descriptions and additional details for traits in the near complete-case dataset.**

| Trait | Description | Data type | Sample size (*n*)* | Units/Levels (%) | Reference |
|---|---|---|---|---|---|
| Activity time | Time of day when species is most active (cathemeral: active at any time (irregular); diurnal: active at day time; nocturnal: active at night time) | Categorical (nominal, multicategorical) | 145 | Cathemeral (11%)<br>Diurnal (58%)<br>Nocturnal (31%) | Meiri (2018) |
| Female snout-vent length | Average female snout-vent length for species | Numerical (continuous) | 137 | Millimetres (mm) | Meiri (2018) |
| Insular endemic | Whether the species lives on an island (Yes) or not (No) | Categorical (nominal, binary) | 152 | No (54%)<br>Yes (46%) | Meiri (2018) |
| Largest clutch | Maximum observed litter size for species | Numerical (count) | 142 | Eggs/Neonates | Meiri (2018) |
| Latitude | Centroid latitude recorded for species | Numerical (continuous) | 152 | Degrees (°) | Roll et al. (2017), obtained from Meiri (2018) dataset |
| Maximum snout-vent length | Maximum observed snout-vent length for species | Numerical (continuous) | 152 | Millimetres (mm) | Meiri (2018) |
| Smallest clutch | Minimum observed litter size for species | Numerical (count) | 141 | Eggs/Neonates | Meiri (2018) |

White and gray rows represent numerical and categorical traits, respectively. * = The number of complete observations in the dataset (up to 10% missingness was permitted for each trait).
