## Supplementary Information file for "A real data-driven simulation strategy to select an imputation method for mixed-type trait data"

#### Consideration of other datasets

In addition to the Meiri [1,2] dataset, trait data from other published datasets were considered for use in this study. These included the amniote life history database [3,4], AnAge [5,6], and vertebrate home range sizes [7,8]. Initially, these datasets were downloaded and merged together in the R environment v. 4.0.3 [9]. The merged dataset comprised observations for 8219 Squamata species and 80 traits. Upon matching to the species names in the multiple sequence alignment of cytochrome *c* oxidase subunit 1 (COI) sequences, many traits were dropped from the dataset as they had less than 100 observations available.

Figure 1 depicts a visualization of the missingness proportion of a subset of these traits (i.e. the 31 traits with the most observations). In Figure 1, the traits that were dropped due to low data availability were egg mass, gestation (days), and incubation (days) from the amniote life history database [3,4]; maximum longevity (captive), maximum longevity (wild), metabolic rate, and temperature from the AnAge dataset [5,6]; locomotion, predator/prey mass ratio, prey mass, thermoregulation, and trophic guild from the vertebrate home range sizes dataset [7,8]; and diet, foraging mode, hatchling/neonate snout-vent length (SVL), largest mean clutch size, and smallest mean clutch size from the Meiri [1,2] dataset.

#### Centroid sequence selection

The first step of the centroid sequence selection process [10] entailed performing a separate multiple sequence alignment for each species. The *AlignTranslation* function from the R package “DECIPHER” v. 2.18.1 [11,12] was used, setting the *geneticCode* argument to “SGC1” for vertebrate mitochondrial DNA. The *dist.dna* function from the R package “ape” v. 5.4.1 [13] was used to construct a genetic distance matrix for each alignment, specifying the TN93 [14] model of DNA evolution. The TN93 model is the most parameter-rich model available in the *dist.dna* function and should sufficiently account for multiple substitutions within a species-level alignment. The centroid sequence was then selected as the sequence in the alignment with the minimum pairwise distance to all other sequences.

#### Phylogenetic eigenvector selection

The functions *Phylo2DirectedGraph*, *PEM.fitSimple*, and *PEM.build* from the "MPSEM" package in R v. 0.3.6 [15] were used to convert the gene trees into phylogenetic eigenvectors. As different trait data were available in each dataset with simulated missingness, a different set of eigenvectors was also required. The method employed in Johnson *et al.* [16] was used to address this task as follows: for a particular gene tree and target trait with simulated missingness, *PEM.fitSimple* was first used to estimate the *a* (steepness parameter) and *psi* (relative evolution rate) parameters using maximum likelihood for those species with trait data available. These parameter values were then specified for the function *PEM.build* to construct a new set of eigenvectors that was expanded to include those species with missing trait data [16]; see [17] for a detailed tutorial on use of these functions). Consequently, in each imputation run, each trait had its own set of eigenvectors that corresponded to each tree.

Phylogenetic eigenvector selection is a topic of contention, and agreement regarding the best practice has not yet been achieved (see [18-19] for reviews of the topic). MPSEM offers utility functions for performing forward stepwise selection to choose a set of eigenvectors. However, use of this function became computationally unfeasible considering the scale of the current study and often resulted in large numbers of eigenvectors (e.g. > 70). An alternative approach used in previous trait imputation studies [20,21] follows that the number of eigenvectors that explain a certain percentage (e.g. 95%) of the variance in the phylogenetic structure are selected for use in the imputation process. In the current study, a threshold of 95% was considered initially (e.g. as used in [20]); however, this threshold often resulted in large numbers of eigenvectors being included in the imputation process (e.g. > 80). As the trait dataset in this study had a limited sample size ($n$ = 152), the use of large numbers of eigenvectors led to unstable results, particularly in the case of *MICE*. To mitigate this issue, the threshold was reduced to 65% (similar percentages were employed successfully in [22]). The number of eigenvectors selected varied according to the tree and the trait under consideration but were more reasonable relative to sample size (i.e. consistently less than 70). As 65% still accounted for a substantial proportion of phylogenetic structure variance, this threshold was deemed suitable for use in the imputation process.

#### Missing at random (MAR) scenarios

To simulate MAR scenarios using real data, a logistic regression model was fitted for each trait using the original Meiri [1,2] dataset. In each model, missingness in the trait of interest was specified as the response variable (1 = observed/0 = missing) and each auxiliary trait was specified as a covariate. A final logistic regression model was fitted that included all traits with significant terms in the univariate models (p-value < 0.05). The final model was used to introduce missing values into the complete-case dataset. It should be noted that some traits had either no missing values in the original dataset (insular endemic) or no significant predictors of missingness upon fitting of the logistic regression models (maximum SVL); consequently, these traits were omitted from the MAR scenarios. For the trait latitude, the logistic regression model was fitted but resulted in an inadequate amount of missingness introduced into the complete-case dataset (less than 0.08). To mitigate this issue, the model's intercept was reduced by half to increase the amount of missingness introduced into the dataset.

#### Missing not at random (MNAR) scenarios

In addition to testing the robustness of the imputation methods in various missingness scenarios, the MNAR simulations were designed to mimic realistic situations in which data may be more likely to be missing for squamates. Values were removed from below the 10$^{th}$ percentile of the data for the traits maximum SVL, female SVL, largest clutch, and smallest clutch. For the trait activity time, 15% of observations from the "nocturnal" category were removed. For the trait environmental numerical trait latitude, values were removed from between 10° and -10° (i.e. near the equator). These choices were made as newly discovered lizard species (i.e. in the 21$^{st}$ century) tend to be smaller in size, nocturnal, and inhabit tropical regions, which suggests that missing data may also skew in these directions [23]. In addition, tropical regions are known to be undersampled, especially in the case of reptiles [23,24]. Finally, 15% of observations from the "yes" category for insular endemic were removed. Although many squamates reside on islands, new species may be less likely to be observed due to their inhabitation of inaccessible or remote locations.

#### Imputation algorithm details and parameter selection

The *kNN* function in the R package "VIM" v. 6.1.0 [25] was used to perform *k*-nearest neighbour (*KNN*) imputation. For a given dataset, distances between variables of different types are calculated (using a Gower distance [26] framework for mixed-type variables) and used to determine the *k* nearest neighbours for a particular variable with a missing observation. The median (default for continuous variables) or most frequent category (default for categorical variables) of the *k* observed values of the nearest neighbours are used to impute the missing value. As results are sensitive to the value of *k* [27], a range of *k* values (1-50) was considered in the current study. For each dataset replication, the value of *k* that resulted in the lowest imputation error rate (mean squared error (MSE) or proportion falsely classified (PFC) for numerical and categorical traits, respectively) was used in the final imputation model.

The “missForest” R package v. 1.4 [28,29] was used to perform random forest (*RF*) imputation. In this method, all missing values are first imputed using the mean or most frequent category for continuous and categorical variables, respectively. To impute values for a particular variable (hereafter referred to as the “target variable”), a random forest is fit to the observed data, specifying the target variable as the response. This process is then repeated in an iterative fashion. The imputed values are updated with each iteration until a stopping criterion is met (i.e. when the differences between the previous and newly imputed values increase). The number of trees (argument *ntree*) grown in the random forests can impact imputation precision [28]. In our study, *ntree* values of 100 and 1000 were considered. For each dataset replication, the *ntree* value that resulted in the lowest imputation error rate (MSE or PFC) was used in the final imputation model.

The final method considered was multivariate imputations using chained equations (*MICE*) (“mice” package v. 3.13.0) [30]. *MICE*, otherwise known as “fully conditional specification”, fits a separate model for each variable using the observed data. The variable to be imputed is specified as the dependent variable, and the other variables in the dataset are specified as the covariates. The fitted model is then used to predict the missing values of the dependent variable. The model used depends on the type of variable being imputed (default = predictive mean matching and logistic regression for continuous and categorical variables, respectively).

*MICE* offers a multitude of other imputation models, including those for numeric, binary, ordinal, and nominal data; as such, the user can select the model that is most appropriate for their data. Akin to the missForest algorithm, this process is repeated iteratively (default number of iterations = 5). As *MICE* is a multiple imputation method, *m* datasets are imputed. Graham *et al.* [31] suggest that larger *m* values can improve statistical power, and Von Hippel [32] recommends that the value used for *m* should correspond to the proportion of missingness in the dataset. Therefore, for each dataset replication in the current study, different values of *m* were considered (5, 10, 40). The value of *m* that resulted in the lowest imputation error rate (MSE or PFC) was used in the final imputation model.

### Phylogenetic signal

All genes exhibited significant levels of phylogenetic signal for the categorical traits activity time and insular endemic (Fig. S1a). As the *D* metric [33] measures the phylogenetic signal of binary traits, the three-level categorical trait activity time was broken down into two indicator variables “activity time: diurnal” and “activity time: nocturnal”, with “cathemeral” being treated as the reference category. For both indicator variables, 1 indicates that this behaviour has been observed in the species, and 0 indicates that no evidence has been found for the behaviour. Except in the case of c-mos for activity time: nocturnal, all genes demonstrated stronger levels of phylogenetic conservation than would be expected under Brownian motion (BM) ($D < 0$) for both activity time: nocturnal and activity time: diurnal. The strongest phylogenetic signal for insular endemic was observed in RAG1, as *D* fell closer to 0 compared to the *D* values for c-mos and COI. All genes exhibited significant levels of phylogenetic signal [34] for all numerical traits (Fig. S1b). For the traits latitude, maximum SVL, and female SVL, $\lambda$ values exceeded 0.80 for all genes, indicative of patterns of trait evolution that more closely adhere to BM. λ values for smallest clutch and largest clutch were also high for COI and c-mos (> 0.90); for RAG1, however, λ values for these traits were comparatively weaker (0.64 and 0.34 for smallest clutch and largest clutch, respectively).
